# Deep Mutational Scanning of the GPCR Rhodopsin Reveals Pleiotropic Mutational Effects and Their Roles in Modulating Light Sensitivity

**DOI:** 10.64898/2026.07.27.740619

**Authors:** Steven K. Chen, Jing Liu, Belinda S.W. Chang

**Author notes:** Correspondence: Belinda S.W. Chang, Ramsay Wright Zoological Laboratories, 25 Harbord Street, Toronto, ON M5S 3G5, Canada. These authors contributed equally (Steven K. Chen, Jing Liu).

## Abstract

Biophysical pleiotropy, the phenomenon in which a mutation affects multiple protein properties, underlies many genetic diseases and shapes protein evolution, yet remains poorly understood, limiting synthetic biology and therapeutic development. Although extensively studied in soluble proteins, little is known about how pleiotropic effects shaped the evolution of receptor protein functions. Here, we developed a cell-based assay designed to assess pleiotropic effects in G protein-coupled receptors (GPCRs), the largest receptor class in eukaryotes. Because our assay design allows for multiple GPCR molecular phenotypes (basal activity, ligand-dependent signaling, and cellular receptor abundance) to be simultaneously assessed, we applied it to study rhodopsin, a visual GPCR that evolved high light sensitivity by maximizing light-driven responses while minimizing thermally driven noise. To investigate this, we integrated our assay with deep mutational scanning of relevant rhodopsin domains to test the impacts of ∼2,000 single-residue substitutions in rhodopsin across all three molecular phenotypes (∼6,000 measurements). By analyzing these data in the context of rhodopsin’s protein structures, we discovered complex pleiotropic effects that act asymmetrically to impose joint constraints, restricting mutational tolerance in rhodopsin’s retinal binding pocket and G protein interaction interface. A dose-response analysis of rhodopsin signaling revealed that these pleiotropic constraints arise from the dual functions of its chromophore, which acts as an agonist upon light-driven isomerization but also an inverse agonist in darkness. Because these constraints reflect mechanistically driven limitations in the evolution of receptor signaling, these findings reveal a role for biophysical pleiotropy in shaping the sensory capabilities of receptor proteins.

**Significance Statement:** Understanding how genetic variation impacts protein function is an important but challenging area of research as many mutations are pleiotropic, simultaneously affecting multiple protein properties, including structure, stability, and activity. Previously, genetic screens have systematically mapped pleiotropic effects in soluble proteins, but transmembrane receptor proteins embedded in cellular membranes are much more difficult to assay. These are key receptors that convert environmental stimuli into appropriate physiological responses. Here, we present a systematic study of pleiotropic effects within the receptor responsible for dim-light vision in vertebrates, revealing how light sensitivity can be mediated through differential conformational states of its chromophore. This is important not only for understanding how receptors evolved enhanced capabilities, but also for the development of tunable protein systems.

## Introduction

Mutations modulate protein functions by exerting diverse, but often unequal, effects on multiple protein properties and activities. These effects can produce highly divergent—even opposing—molecular phenotypes, leading to trade-offs between different aspects of protein functions (1–3). For example, a mutation that increases a protein’s activity can simultaneously reduce its stability, thereby lowering the total amount of functional protein within a cell (1, 4). Such relationships between mutations and their multiple effects, termed biophysical pleiotropy, constrain evolutionary pathways and define the extent to which proteins tolerate random genetic changes in their coding sequences (1, 4, 5). Consequently, the pleiotropic effects of mutations have important implications for many areas of biology, including the evolution and regulation of protein functions, signal transduction, and homeostatic control. Mutations that disrupt these processes are often associated with human genetic diseases (3, 6, 7), and when they occur in protein drug targets, can cause adverse drug responses and side effects (8–11). Therefore, understanding the pleiotropic effects of mutations is of fundamental importance for addressing modern challenges in evolutionary biology, synthetic biology, physiology, and medicine.

In protein evolution, pervasive biophysical pleiotropy imposes severe constraints on the number of selectively accessible evolutionary pathways (12). Whether a mutation contributes to the evolution of a new or enhanced protein function is contingent not only on the genetic background in which the mutation occurs (that is, epistasis), but also on its effects on multiple protein properties (that is, biophysical pleiotropy). This understanding of protein evolution is supported by studies that compared pleiotropic effects on protein stability with catalysis (13–17), antibiotic resistance (12, 13), dynamics (18–21), and binding (22–25). Yet, this concept and other related principles of protein evolvability are largely derived from studies of soluble proteins, which are characterized by well-defined binding site(s), water-exposed surface residues, and a hydrophobic core. In contrast to soluble proteins, comparatively little is known about how pleiotropic effects shaped the evolution of receptor protein functions.

To address this knowledge gap, we selected the G protein-coupled receptor (GPCR) rhodopsin, a photoactivated transmembrane receptor responsible for dim-light vision in vertebrates. Covalently bound to its 11*-cis-*retinal chromophore absorption of a photon triggers isomerization to the all*-trans* configuration, leading to formation of the biologically active metarhodopsin II form. These conformational changes activate the heterotrimeric G protein transducin, which initiates a signaling cascade and ultimately results in vision. Rhodopsin is an ideal system for dissecting biophysical pleiotropy because its function depends on the simultaneous optimization of multiple biophysical properties. Rhodopsin is known to exhibit extremely low thermal noise, which is indistinguishable from light-driven activation, but when paired with its quantum efficiency of nearly 70% (26), produces an extremely high light sensitivity. It has thus been proposed that rhodopsin must have evolved optimizations, potentially through pleiotropic mutations, to maximize light-driven responses while minimizing thermally driven noise (27, 28).

To investigate how pleiotropic effects might have shaped the evolution of receptor-mediated signaling, we turned to deep mutational scanning (DMS), a high-throughput reverse genetics approach capable of resolving the effects of thousands of genetic mutations (29, 30). Multiphenotypic DMS that measure multiple protein properties are especially powerful for resolving complex pleiotropic effects. However, most DMS studies of GPCRs have focused on a single molecular phenotype (29, 31–35). In the case of rhodopsin, mutational scans have been limited to quantifying cell surface expression (31–34) since developing a high-throughput screen of its native function (phototransduction) is especially challenging.

Here, we overcome these challenges by developing a cell-based system capable of measuring three aspects of rhodopsin function: basal (dark) activity, light-dependent signaling, and cellular receptor abundance. We integrated our assay with multiphenotypic DMS to measure, for each of these molecular phenotypes, the impacts of nearly all possible single amino acid substitutions (∼2,000 variants) across four rhodopsin domains. These domains—comprising extracellular loop 2 (EL2), transmembrane helix 5 (TM5), intracellular loop 3 (ICL3), and TM6—are thought to be key for rhodopsin function because they undergo large structural rearrangements during photoactivation (36, 37), enabling allosteric signal transmission from rhodopsin’s ligand binding site to its cytosolic surface (38, 39). The resulting atlas of variant effects serves not only as a dataset for studying the pleiotropic effects of mutations in *RHO* but also a resource for interpreting *RHO* missense variants of uncertain significance relevant to visual disease (40). By analyzing these data in the context of rhodopsin’s protein structures, we discovered that residues located within rhodopsin’s retinal binding pocket (RBP) and G protein interaction interface (GPI) display strong asymmetries in their pleiotropic effects. Through an extensive comparison of rhodopsin signaling before and after light-driven isomerization of its chromophore, we revealed the molecular basis of these pleiotropic constraints.

## Results

### Cell-Based Assay of Rhodopsin Dark Activity, Light Activity, and Cellular Abundance

To investigate the pleiotropic effects of mutations in *RHO*, we developed a two-color system in the yeast *Saccharomyces cerevisiae* capable of simultaneously monitoring the activity and cellular abundance of rhodopsin variants in response to treatment with retinal and light **(Fig. 1*A*, and *SI Appendix,* Fig. S1)**. This system also allows us to measure rhodopsin activity in the dark, which is a measurement of the activity of rhodopsin bound to the retinal chromophore, without light activation. For this, we leveraged an assay that we previously described, which uses a chimeric Gα protein to couple rhodopsin activity to a modified GPCR signaling pathway in yeast (41). We further modified this system to incorporate two additional aspects of rhodopsin function. First, we engineered a pathway-responsive eGFP fluorescent reporter as a readout of rhodopsin-mediating signaling. Second, we fused rhodopsin to the fluorescent protein ymScarlet (RHO-ymS) to monitor its cellular abundance. Because this system enables multiple rhodopsin molecular phenotypes to be simultaneously measured, it should allow certain biophysical ambiguities to be resolved such as determining whether increases in receptor signaling reflects a true increase in light-dependent activation or instead arises from elevated basal activity.

**Figure 1.**
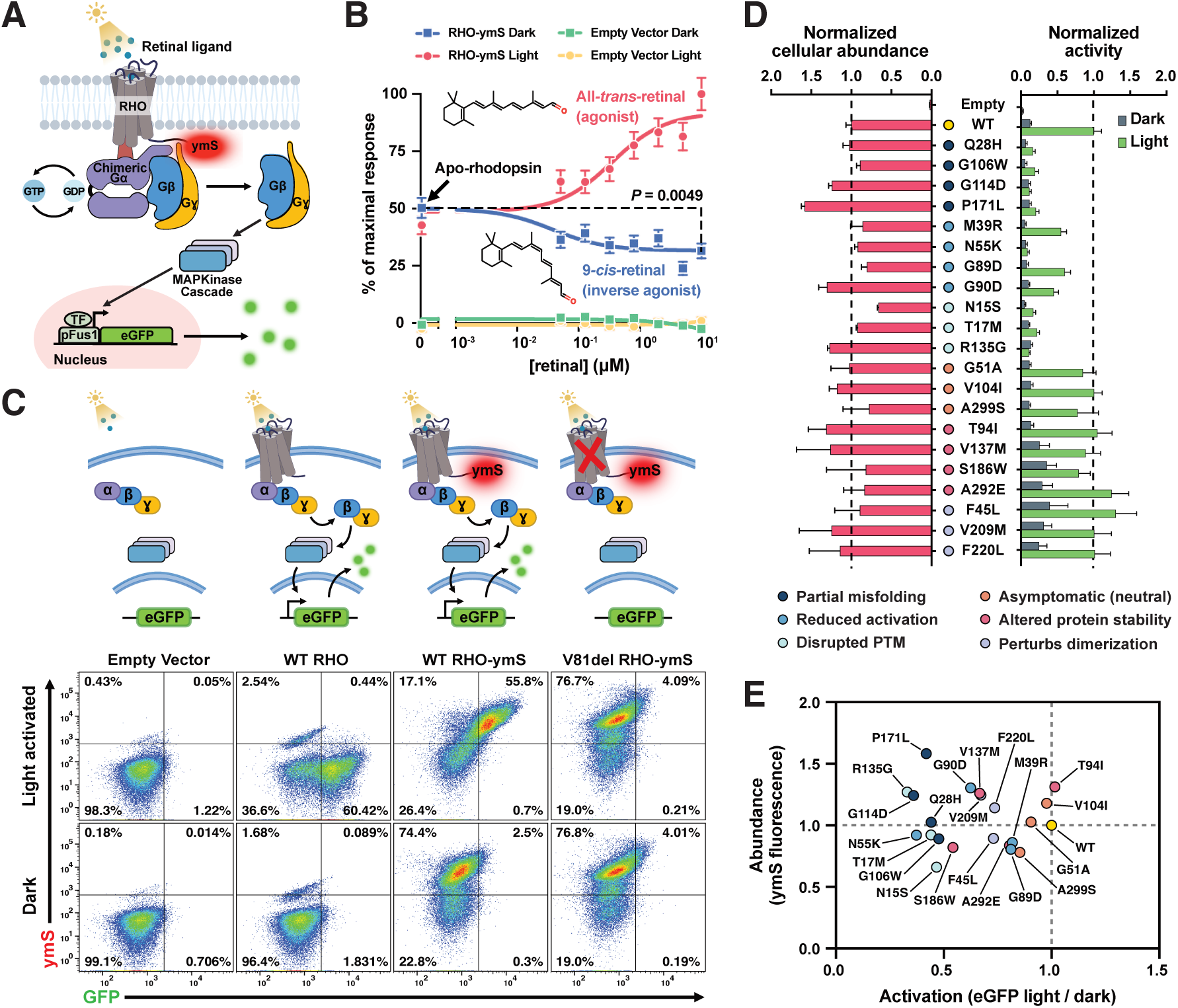
Yeast-based biosensor of rhodopsin basal (dark) activity, light activation, and cellular abundance. (*A*) Schematic showing light-dependent rhodopsin activation of an engineered GPCR signaling pathway in yeast. (*B*) Rhodopsin-mediated signaling as a function of retinal concentration. Comparison of the 0 and 10 μM conditions of retinal confirmed the inverse agonist effect (*P* = 0.0049). Data are mean ± s.d. of *n* = 4 biological replicates for RHO-ymS and *n* = 2 for empty vector controls. Statistical analysis was performed using unpaired two-tailed Student’s t-test. (*C*) Flow cytometric measurements of rhodopsin activity as a function of eGFP fluorescence and cellular rhodopsin abundance as a function of ymS fluorescence under dark and light conditions. (*D*) Rhodopsin activity and cellular rhodopsin abundance measurements for 21 missense rhodopsin variants. Data are mean ± s.d. of *n* = 4 biological replicates. (*E*) Comparison of rhodopsin activation and cellular abundance using the data from (*D*).

To evaluate the performance characteristics of our system, we first expressed RHO-ymS in the engineered yeast and assessed rhodopsin-mediated signaling in response to treatment with retinal and light. A dose-response experiment, in which rhodopsin-expressing yeast cells were incubated with the rhodopsin ligand analog 9*-cis-*retinal followed by irradiation with white light, showed a dose-dependent increase in signaling activity with retinal concentration **(Fig. 1*B*)**. No increase in signaling was observed in rhodopsin-expressing yeast in the dark or in empty vector controls, demonstrating that pathway activation is driven by rhodopsin and requires both retinal and light. Notably, rhodopsin in the apo state (apo-rhodopsin), lacking the retinal chromophore, exhibited elevated basal activity compared to the empty vector controls. This basal activity, likely arising from thermally induced light-independent receptor activation, was partially suppressed when apo-rhodopsin was reconstituted with 9*-cis-*retinal, consistent with the understanding that retinal in the *cis* conformation acts as an inverse agonist against vertebrate opsins (42). Of physiological importance, this interaction between *cis* retinal and rhodopsin has been shown to account for the extremely low thermal activation rate of dark-adapted rhodopsin, preventing noise that would otherwise interfere with real-light detection in photoreceptors (43–47).

Having demonstrated that our system is capable of producing physiologically relevant readouts of both dark- and light-activated rhodopsin activity, we next assessed steady-state cellular rhodopsin abundance to distinguish variants that perturb rhodopsin activity directly from those that act by altering cellular rhodopsin abundance. For this, we compared RHO-ymS to the patient-derived loss-of-function rhodopsin mutant V81del (48) fused to ymScarlet (V81del RHO-ymS). Flow cytometric analysis of light-activated RHO-ymS-expressing yeast cells showed a predominantly ymS^+^eGFP^+^ population, whereas cells maintained in the dark were ymS^+^eGFP^-^ **(Fig. 1*C*)**. In contrast, the V81del variant remained ymS^+^eGFP^-^ regardless of light activation, suggesting that this loss-of-function variant compromises light-dependent rhodopsin-mediated signaling through mechanisms independent of changes in cellular receptor abundance.

To further validate our system, we expanded the scope of our assessment to include a panel of 21 single amino acid substitution *RHO* variants spanning six categories. These include variants previously characterized *in vitro* to cause partial misfolding (49–51), reduced light activation (52–55), disrupted post-translational modification (PTM) (56–58), altered receptor stability (51, 59–63), perturbed receptor dimerization (64–66), or are phenotypically neutral (67–72). Our measurements of dark- and light-dependent activities are broadly concordant with the known molecular phenotypes of these variants, whereas substantial variation was observed in the abundance assay **(Fig. 1*D* and *SI Appendix,* Fig. S2)**. Many variants that cause partial misfolding or disrupted PTM showed large reductions in rhodopsin activation but mixed effects on cellular abundance, suggesting that these variants act primarily by attenuating rhodopsin activation. Consistent with this observation, variant abundance did not correlate with activation **(Fig. 1*E*; Pearson’s r = -0.09)**, indicating that these two molecular phenotypes contribute orthogonally to the net mutational effect of each protein variant. Notably, we also found that many variants characterized to alter rhodopsin stability or perturb its dimerization retained light-dependent signaling activity comparable to, or even exceeding, that of wild-type **(Fig. 1*D*)**. However, these variants also showed elevated activity in the dark, suggesting a potential trade-off limiting further enhancements in signal-over-noise ratios. Together, these results establish our engineered system as a platform capable of dissecting multiple physiologically relevant rhodopsin molecular phenotypes, including its dark activity, light-dependent signaling activity, and steady-state cellular abundance.

### Sequence-Function Maps of Rhodopsin Revealed by Multiphenotypic Deep Mutational Scanning

The agonist and inverse agonist effects of retinal on rhodopsin activity (**Fig. 1*B***) suggest that rhodopsin is evolutionarily optimized to maximize light-driven responses while minimizing thermally driven noise. To investigate the potential role of pleiotropic mutational effects in shaping these optimizations, we combined our multiphenotypic cell-based assay with DMS. For this we synthesized a scanning site-saturation mutation library that encodes all possible single amino acid substitutions for sites 177–277 in rhodopsin starting from the beginning of EL2 to the end of TM6. EL2–TM6 is a region of functional importance and likely contain residues involved in optimizing the range of rhodopsin activation because: (i) EL2, containing two antiparallel β-sheets (β3-β4), forms a ‘plug’ over the retinal binding pocket which helps to stabilize the retinal chromophore (73, 74); (ii) during rhodopsin activation, TM5 and TM6 has been shown to facilitate anisotropic signal propagation from rhodopsin’s RBP to its GPI (38, 39); (iii) ICL3, the region connecting TM5 and TM6, forms critical interactions with downstream G proteins during rhodopsin-mediated signaling (75–77); and (iv) ICL3 undergo very large conformational changes during receptor activation but is predicted to tolerate amino acid substitutions without major stability loss **(Fig. 2*A*,*B* and *SI Appendix,* Fig. S3)**. In total, the mutated region encodes a theoretical maximal diversity of 2,121 single amino acid substitution variants, including variants encoded by premature termination codons.

**Figure 2.**
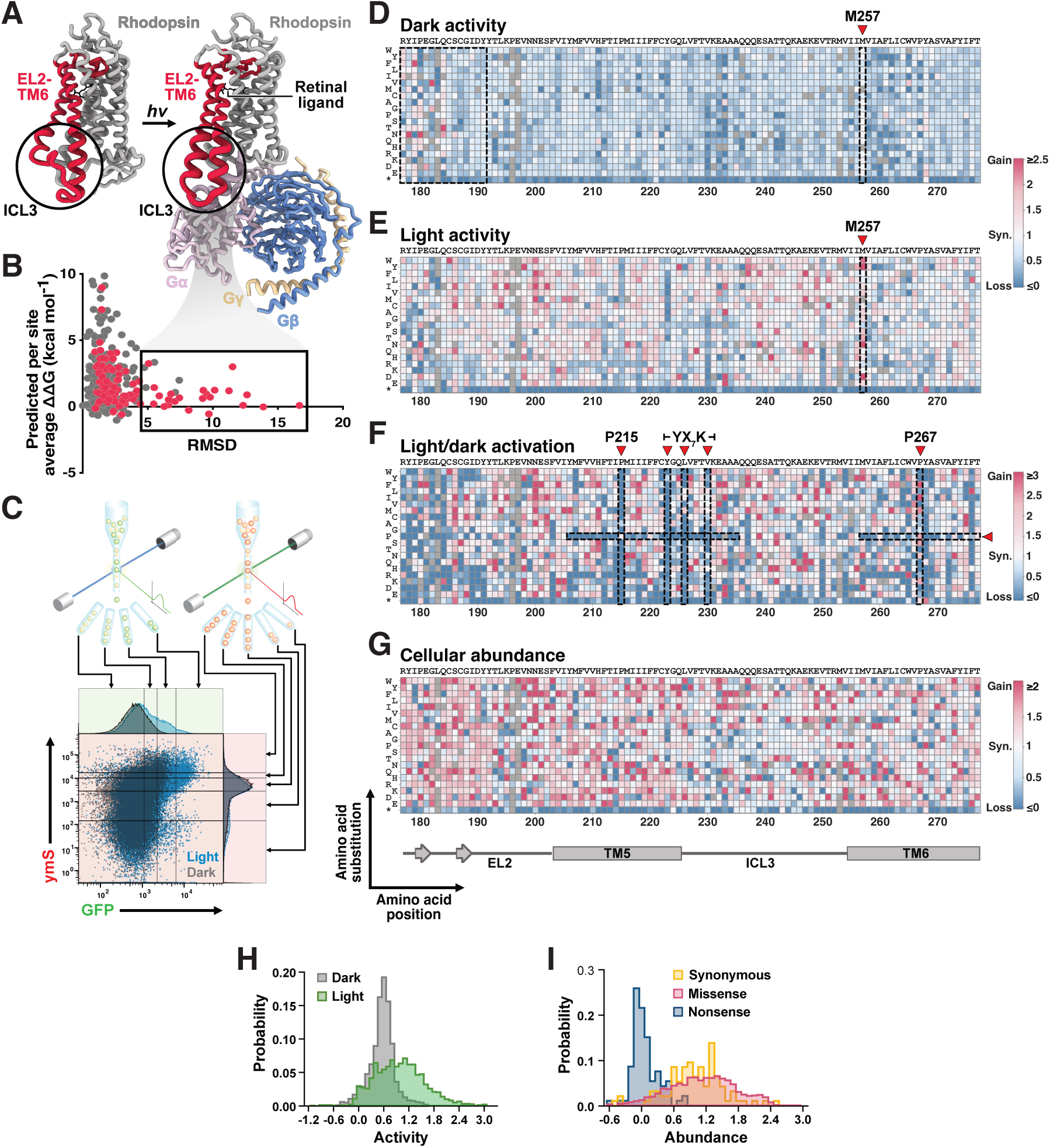
Mapping the effects of rhodopsin variants across multiple molecular phenotypes. (*A*) Protein structures of rhodopsin in the dark state (PDB: 1U19) and in the active state in complex with heterotrimeric G proteins (PDB: 6OY9). Extracellular loop 2 to transmembrane domain TM6 (EL2-TM6) are colored crimson. (*B*) Across sites comparison of predicted protein stability changes upon amino acid replacement versus the root-mean-square deviations (RMSD) in α-Carbon atomic positions between dark and active state rhodopsin structures. Protein stability change data represent the mean FoldX predicted Gibbs free energy change for all possible amino acid replacements at each position in rhodopsin. (*C*) Fluorescence-activated cell sorting strategy for sorting rhodopsin variants based on their signaling activities (eGFP) and cellular abundances (ymS). (*D-G*) Variant effect maps of rhodopsin dark activity, activity when irradiated with white light (light activity), light-over-dark activation, and cellular abundance. Scores are represented with colors from low (blue) to high (red) with missing variants in grey. Data was scaled by centering premature termination variants to 0 and synonymous variants (Syn.) to 1, with the exception of the dark activity map, which was scaled according to the light activity map to enable direct comparisons. (*H*) Distributions showing variant effects on rhodopsin activity under dark (grey) versus light (green) conditions. (*I*) Distributions showing variant effects on cellular rhodopsin abundance.

To resolve pleiotropic effects on rhodopsin dark activity, light activation, and cellular receptor abundance, we applied 2-dimensional fluorescence-activated cell sorting coupled to deep sequencing (2D FACS-Seq). This involved expressing the library in the engineered yeast such that each cell expresses a single variant. The variants are then sorted in three independent experiments: (i) based on eGFP to assess variant basal activity in the dark; (ii) based on eGFP to assess variant activity when exposed to light; and (iii) based on ymScarlet to assess variant abundance **(Fig. 2*C* *and* *SI Appendix* Fig. S4)**. After FACS, variants were extracted from the sorted cells, deep sequenced, and analyzed in the context of fluorescence intensity distributions acquired during FACS **(*SI Appendix,* Fig. S5)**. Through this approach, we generated variant effect maps that capture the dark activity, activity when irradiated with white light (light activity), light-over-dark activation, and cellular abundance of 2,016 rhodopsin variants **(Fig. 2*D*-*G*)**. These maps provide variant effect scores for 95.0% of all possible single amino acid substitutions and premature termination variants within EL2–TM6 for each of the measured molecular phenotypes.

To assess the quality of our variant effect scores, we first examined variant effect distributions. As expected, variants kept in the dark were annotated with low activity scores, forming a narrow distribution at the bottom end of the scale **(Fig. 2*H*)**. By contrast, many variants exposed to light were annotated with higher scores, indicating that a substantial proportion of the variants can be activated by light. Across all datasets, variants that contain premature termination codons feature very low scores with distributions that segregate from those encoded by synonymous mutations **(Fig. 2*I* and *SI Appendix,* Fig. S6)**.

### Sequence-Function Map in the Context of Rhodopsin Biochemistry

We also evaluated our variant effect maps against known rhodopsin biochemistry. Amino acid substitutions of residues proximal to the β3-β4 hairpin in EL2, thought to stabilize the chromophore and suppress spontaneous receptor activation in the dark (73, 74), show elevated dark activity **(Fig. 2*D*)**. Substitutions of residue M257 also show higher dark activity, which is consistent with previous studies that have found amino acid replacements at this site to lower the energy barrier between rhodopsin’s inactive and active states (78) **(Fig. 2*D*)**. In particular, the M257Y mutant, which has been characterized in depth to increase dark activity by ∼40% and further increase activity when reconstituted with the rhodopsin agonist all*-trans-*retinal (79), displays remarkably similar trends in our maps **(Fig. 2*D*,*E*)**. Disruptions to evolutionarily conserved prolines (Pro215 and Pro267), responsible for facilitating outward motions and rotations of TM5 and TM6 during receptor activation (80, 81), severely compromise rhodopsin activation **(Fig. 2*F*)**. In contrast, introducing prolines—a potent helix breaker (82)—at most other positions within the TM domains renders rhodopsin incapable of being activated by light. Substitutions of ‘hydrophobic patch’ residues (Tyr223, Leu226, and Val230) within the YX_7_K motif, a highly conserved ‘microswitch’ conserved across class A GPCRs thought to stabilize the receptor’s active state and form coupling interactions with the heterotrimeric G proteins (83–86), are also poorly tolerated **(Fig. 2*F*)**. These results are all consistent with known rhodopsin biochemistry, and together with our analysis of variant effect distributions, support the quality of our dataset.

### Asymmetries in Pleiotropic Mutational Effects

To identify residues in rhodopsin involved in maximizing light-driven signals while minimizing thermally driven noise, we calculated median scores across mutated sites and visualized them in the context of rhodopsin’s protein structures **(Fig. 3*A*,*B*)**. Median scores of variant dark activities appear to be generally homogenous across sites while light activities, activation, and cellular abundances show substantially more variation. However, median scores of variant activation are visibly lower in the RBP, suggesting that amino acid substitutions of residues involved in binding the retinal chromophore severely compromise light-dependent receptor activation **(Fig. 3*B*,*C*)**. Similarly, substitutions in the GPI, regions on the cytoplasmic face of the protein that interact with the G protein transducin during receptor-mediated signaling (85, 75–77), also appear to lower activation **(Fig. 3*B*,*D*)**. We therefore prioritized analyzing RBP and GPI residues. Substitutions of RBP residues showed large reductions in median light activities and activation, but also significant increases in dark activities and cellular receptor abundances **(Fig. 3*E* and *SI Appendix,* Fig. S7)**. In contrast, substitutions of GPI residues showed large reductions in light activities and light-dependent activation but limited effects on basal activities and abundances.

**Figure 3.**
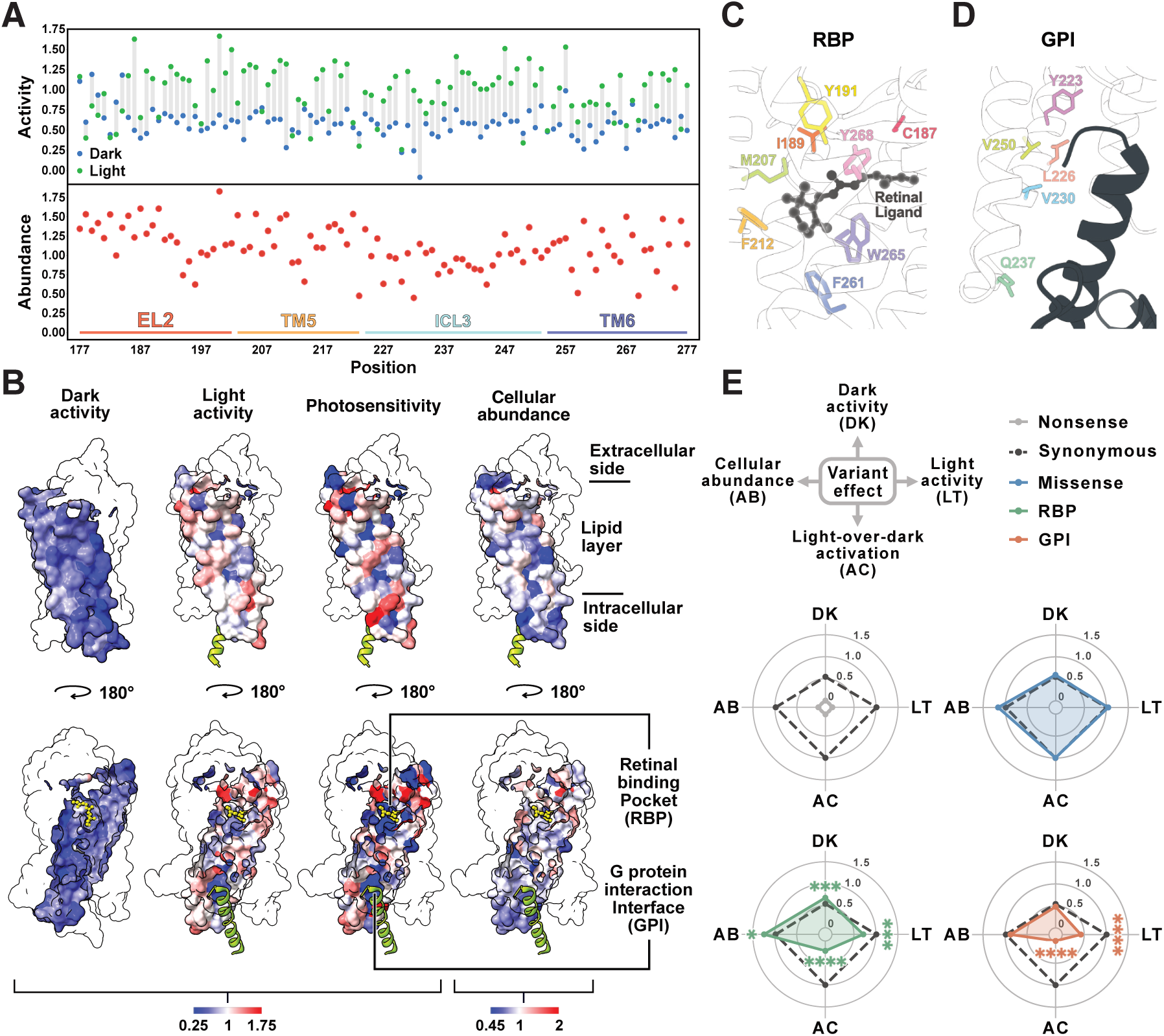
Pleiotropic mutational effects in the context of rhodopsin’s protein structures. (*A*) Median variant effect scores across sites in EL2–TM6. (*B*) Median variant effect scores mapped onto rhodopsin protein structures. (*C*) Close-up view of the RBP with the retinal ligand shown in black and residues within 3.7 Å from retinal in EL2–TM6 in color. (*D*) Close-up view of the GPI. The C-terminal α5 Helix of the G protein transducin is shown in black. Key residues on the rhodopsin-side of the interface are shown in color. (*E*) Analysis of variant dark activity, light activity, activation, and cellular abundance for all single amino acid substitutions in EL2–TM6 at RBP and GPI sites. Significance was determined using Kruskal-Wallis test with Benjamini-Hochberg correction (\**P* < 0.05, \*\*\**P* < 0.001, \*\*\*\**P* < 0.0001).

These differences in median values of mutational effects are also reflected in the individual variant scores of the RBP and GPI domains. We examined the relationship of effects on light and dark rhodopsin activities by analyzing the overall direction and magnitude of differences with respect to overall scores **(Fig. 4*A*)**. 42.7% of variants at RBP sites were found to be overrepresented relative to all other sites in the upper left quadrant, associated with decreased light activity and increased basal activity **(Fig. 4*B*)**. In contrast, 56.6% of variants at GPI sites tended to cluster in the lower left quadrant toward decreases in both light and basal activity **(Fig. 4*C*)**, trends which are reflected in the overall magnitude and direction of the differences in scores in the RBP and GPI domains **(Fig. 4*D*)**.

**Figure 4.**
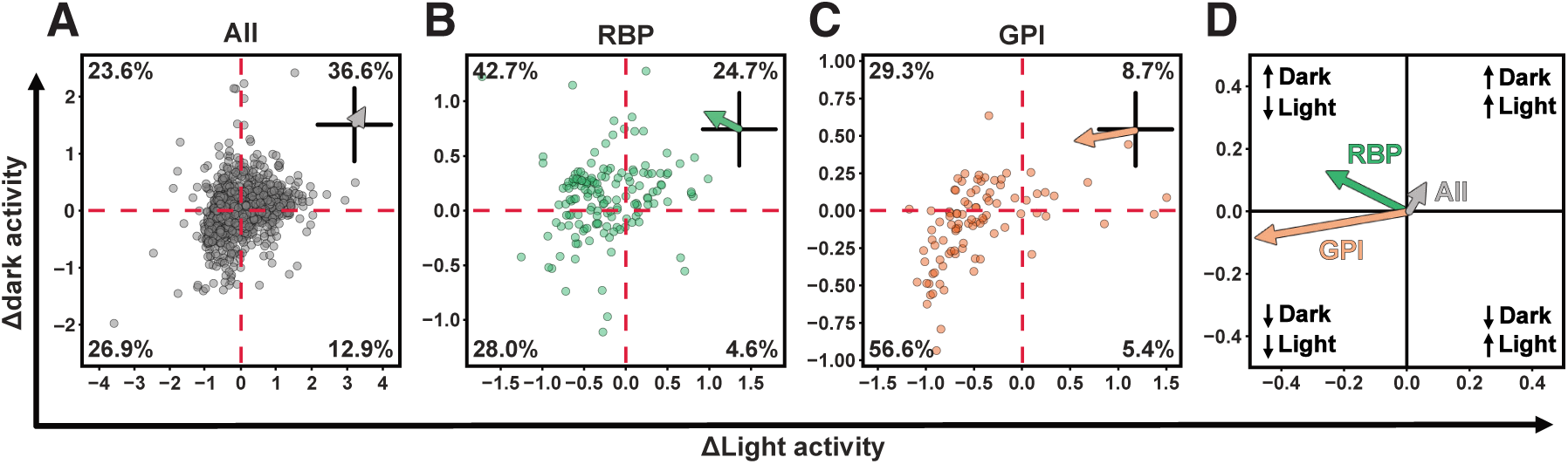
Relative effects of amino acid substitutions at RBP and GPI sites on rhodopsin light and dark activities. (*A*) The effects of amino acid substitutions across all mutated sites (All) compared to the median effects of synonymous variants (red dashed lines). (*B*) As in (*A*) but for substitutions at RBP sites. (*C*) As in (*A*) but for substitutions at GPI sites. (*D*) The colored arrows represent the magnitude and direction of change from the origin to the centroid of each cluster from (*A-C*).

Mechanistically, these asymmetries in pleiotropic effects can be reconciled with our current understanding of the distinct functional roles of each protein region. Because the retinal chromophore functions as an inverse agonist when bound to rhodopsin, substitutions of RBP residues perturb rhodopsin-retinal interactions, leading to increased spontaneous receptor activation and elevated signaling in darkness. Concurrently, these same substitutions negatively impact light-driven rhodopsin signaling activity because proper binding of retinal to rhodopsin is a prerequisite in forming the visual pigment required for real-light detection. Moreover, given that the loss in activation is accompanied by a concomitant increase in steady-state receptor abundance, this trade-off is in support of the hypothesis that residues involved in ligand-binding are not optimized for stability (14). In contrast to the RBP, substitutions of GPI residues tend to only reduce light-dependent rhodopsin activation, suggesting that the GPI is not functionally involved in suppressing thermally driven noise but is rather highly specialized in its role of transducing signals from the receptor to its downstream effector proteins.

## Discussion

Mutations are generally assumed to be pleiotropic at the biochemical level—simultaneously impacting protein activity, stability, aggregation, and interactions—albeit to varying degrees (1–3). Because the effects of individual mutations differ in the extent to which they affect each of these protein properties, single amino acid substitutions that improve one aspect of protein function may be detrimental to another. Therefore, understanding how these effects differ within the context of a protein’s tertiary structure requires assessing the impacts of many single amino acid substitutions on multiple molecular phenotypes and across multiple protein sites.

One of the most interesting results of our deep mutational scanning investigation is the opposing effects of mutations in the retinal binding pocket on key aspects of rhodopsin function. As the first step in the visual transduction cascade of the photoreceptors of the eye, multiple aspects of rhodopsin function are thought to be operating at biophysical limits of the system. This is particularly true with respect to the isomerization of the 11-*cis*-retinal chromophore, the first critical step in formation of the biologically active metarhodopsin II form. Photon absorption by the chromophore, resulting in isomerization to the all*-trans* form, is known to be one of the fastest, most efficient processes in the natural world (87). At the same time, rhodopsin covalently bound to the 11-*cis*-retinal chromophore in the dark exhibits a high barrier to thermal isomerization (88). This means that rhodopsin appears to be optimized not only to maximize light-driven activation, but also for suppression of thermally driven signals, which would otherwise contribute to noise (89). In our deep mutational scanning experiments, we were able to demonstrate that amino acid substitutions in the retinal binding pocket compromised light-dependent activation, while increasing dark activity. This is exactly what would be predicted based on the evolution of rhodopsin, to maximize light response while minimizing thermally driven noise.

Our results also show that RBP and GPI residues display significant differences in their pleiotropic effects. Although the two protein regions have been shown to be connected via an allosteric network responsible for propagating light-driven signals from the RBP to the GPI (38, 39), our analysis show that they exhibit distinct pleiotropic profiles. Amino acid substitutions in both protein regions compromise rhodopsin activation but exhibit different effects on dark activity and cellular abundance. Substitutions of RBP residues tend to decrease receptor activation while increasing basal activity and cellular abundance. This finding is consistent with the activity-stability hypothesis, which proposes that residues involved in ligand-binding or catalysis are optimized for activity rather than protein folding/stability (14). Previous studies that have investigated amino acid replacements in the catalytic site of globular enzymes have generally found support for the activity-stability hypothesis (13, 14, 90–92), for which we now also provide evidence in a GPCR. In contrast to the RBP, substitutions of GPI residues tend to only impair receptor activation without significant effects on dark activity or cellular abundance, suggesting that the trade-off between activity and stability does not extend to the rhodopsin-G protein interaction interface. We note that the aspects of protein stability discussed here, related to protein folding free energies, is different from our measurements of rhodopsin basal activity, which is instead related to the thermal activation of the chromophore in darkness (89).

There are potential caveats that should be considered when interpreting these results, particularly with respect to activity-stability trade-offs, which rest on the assumption that variant abundance is correlated with stability. Although cellular abundance is often used as a proxy for stability in DMS studies (93, 94), it presumes that destabilizing amino acid substitutions cause protein misfolding, resulting in accelerated clearance of misfolded variants by protein quality control mechanisms (95, 96). However, in certain cases this may not be true (97, 98), and more detailed assessments of variant effects on protein aggregation may be useful, potentially through future applications such as high-speed fluorescence image-enabled cell sorting (99).

Our investigation into the pleiotropic effects of mutations in *RHO* provides a window into its evolutionary history. Our exhaustive mutagenesis experiments provide support for the notion that rhodopsin is at an evolutionary peak in terms of not only light activation, but also suppression of thermal noise (88, 100). These fascinating ideas have garnered much support, but until the development of high-throughput assays of rhodopsin activation, investigations mainly centered on comparative studies of visual pigments (89). Our exhaustive mutagenesis experiments provide an independent line of evidence supporting this hypothesis and point the way towards future studies of rhodopsin evolution.

Our investigations are also relevant to transmembrane receptors in general. Our observations of asymmetry in pleiotropic effects—between residues involved in ligand-binding and those that form rhodopsin-G protein interaction interfaces—may represent a general feature of missense mutations in GPCRs and, more broadly, other transmembrane receptor proteins. While further studies will be required to fully validate these findings, the approaches and methodologies demonstrated here provide a solid foundation for investigating these principles in other GPCRs, ultimately enabling more comprehensive mapping of mutational effects across this critical protein superfamily.

## Materials and Methods

### Yeast Strains and Culture

To create a cell-based system capable of simultaneously measuring rhodopsin activity and cellular abundance, we engineered the yeast strain JL701, with genotype W303 MATa far1D mfa2D::pFUS1-eGFP-his3 Sst2D::HygB Ste2D::Trp1 Gpa1D::Gpa1(DCGLF)-Leu2, from an ancestral strain that we previously reported (48). All genome editing steps were performed using a standard LiAc/SS carrier DNA/PEG heat shock method (101) to facilitate knock-ins and knock-outs of selectable markers through homologous recombination. In all cases, integration of DNA cassettes by homologous recombination made use of 800 bp flanking homology arms identical to the sequences flanking the target reading frame. For auxotrophic selection, transformed yeast were plated on synthetic defined (SD) media minus the appropriate amino acid: 6.74 g l^-1^ yeast nitrogen base without amino acids (Bioshop Canada, Cat#YNB406), 20 g l^-1^ glucose, 20 g l^-1^ agar (Bioshop Canada, Cat#AGR001), and 2 g l^-1^ synthetic drop-out media minus the appropriate amino acid. For antibiotic selection, transformed yeast were plated on YPD plus the appropriate antibiotic: 65 g l^-1^ YPD agar (Bioshop Canada, Cat#YPD001) containing Hygromycin B (Wisent, Cat#450-141 XL) at a final concentration of 200 μg ml^-1^ or G418 (Wisent, Cat#400-141 IG) at a final concentration of 500 μg ml^-1^. The final JL701 strain and all intermediate strains were confirmed by counterselection, PCR, Sanger sequencing, and by functional tests using flow cytometry and fluorescence microscopy **(*SI Appendix* Fig. S8)**.

### Plasmid Construction

Plasmids were constructed using standard cloning methods, including Golden Gate and Gibson Assembly (102). Yeast codon optimized mScarlet (ymScarlet) (103) was PCR amplified from a plasmid obtained from Addgene (Addgene Plasmid #111917) and subcloned into a pRS316 expression vector containing the constitutively active GPD promotor, the human *RHO* gene (NM_000539.3), and an *Adh* terminator. A sequence that encodes a flexible 9-amino acid glycine linker with the sequence GGGGSGGGG was synthesized by Integrated DNA Technologies and inserted between the human *RHO* C-terminus and ymScarlet to enable expression of human rhodopsin fused to ymScarlet. All mutations in *RHO* were introduced using partially overlapping primer pairs (104). The cloning vector containing the DNA cassette for integrating the mating pathway-responsive eGFP reporter was assembled from the selection marker his3, the FUS1 promoter (defined as the sequence 1,636 bp upstream of the FUS1 start codon), eGFP (105), and flanking homology arms comprising 800 bp upstream of the MFA2 start codon and 800 bp downstream of the MFA2 termination codon. All other DNA cassettes for yeast genome editing were designed in the same way, with 800 bp flanking homology arms PCR amplified from *S. cerevisiae* Genomic DNA (Millipore Sigma, Cat#69240-3), and either a nutrient or antibiotic selection marker.

### Cell-based Assay of Rhodopsin Dark Activity and Light-Dependent Activation

Cells were inoculated into the appropriate culture media and grown overnight for 18 h in a shaking incubator set to 30°C and 225 rpm. After 18 h, the overnight cultures were refreshed to an OD_600_ of 0.1 (corresponding to ∼620,000 cells ml^-1^) in 50 ml LiteSafe Light Blocking Centrifuge Tubes (Cole-Parmer, Cat#06344-60) or 2.2 ml Deep Well Plates (Luna Nanotech, Cat#503002) wrapped in two layers of tin foil with a tin foil lid. The refreshed cells were then transferred to a dark room, dosed with 9-*cis*-retinal (Millipore Sigma, Cat#R5754) to a final concentration of 10 μM, and then grown for 3 h in a shaking incubator set to 30°C and 225 rpm (50 ml tubes) or 300 rpm (deep well plates). After 3 h, cells were light activated for 3 min under white light using a Fiber-Lite 150w Halogen light source (Dolan-Jenner, Cat#Mi-150) equipped with EEG28 Series Dual Arm Gooseneck Fiber Optic Assembly (Dolan-Jenner Industries, Cat#EEG2823M), transferred to a dark room, dosed with 9-*cis*-retinal to a final concentration of 10 μM, and transferred back to the shaking incubator for 55 min. The 3 min light exposure, 9-*cis*-retinal addition, and 55 min shaking incubation was repeated 6x. The same procedure, without light exposure, was used to measure rhodopsin activity in the dark. Dose-response light (and dark) activation experiments were performed using the same procedure, except the concentration of 9-*cis*-retinal was adjusted depending on the sample. The experimental conditions described here were optimized through a time-resolved dose-response experiment **(*SI Appendix* Fig. S9)**.

### Flow Cytometry

Cultured cells were passed through 35 μm filter caps into 5 ml Falcon tubes (Falcon, Cat#0877123) or 20 μm pore size mesh nylon membranes into Multiscreen 96-well plates (Millipore Sigma, Cat#MANMN2010). Cycloheximide (Millipore Sigma, Cat#01810) was then added to a final concentration of 10 μg ml^-1^ to freeze protein synthesis. In the case of photoactivation experiments, cells were incubated for 10 min at 25°C in a dark room after adding Cycloheximide. The tubes or plates containing the cells were subsequently transferred onto ice in preparation for analysis by flow cytometry. Flow cytometry was performed using a BD FACSymphony A3 Cell Analyzer equipped with a High Throughput Sampler (BD Biosciences) and the BD FACSDiva software (BD Biosciences, v9.3). An example of the gating hierarchy used for all flow cytometry experiments is provided **(*SI Appendix* Fig. S10)**.

### In Silico Mutational Scan and Root Mean Square Deviations

FoldX position scan (v5.1) (106) and PoPMuSiC (v2.1) (107) were used to predict the Gibbs free energy change for all possible single amino acid substitutions in rhodopsin. Rhodopsin in the inactive (PDB: 1U19) and active (PDB: 3PQR) states were used as the starting structures for the predictions. RMSDs in α-carbon atomic positions between the two structures were calculated using UCSF ChimeraX (v1.5) (108).

### Site-Saturation Mutagenesis Library

The site-saturation mutagenesis library, comprising full-length linear double stranded DNA *RHO* variants, was synthesized by Ranomics using the VariantFind platform. *RHO* variants were amplified using KOD Xtreme polymerase kit (Millipore Sigma, Cat#71975) in a touchdown PCR reaction (***SI Appendix,* Table S1,2**) to add flanking AarI restriction enzyme cut sites for scarless integration of the *RHO* variant sequences into a pRS316 expression vector containing the GPD promoter and ymScarlet fused in-frame to a 9 amino acid glycine linker sequence at its 5’ end. After PCR, the PCR products were digested with 20 U of DpnI (New England Biolabs, Cat#R0176) for 3 h at 37°C, checked for a single band using a standard 1% TAE agarose gel, and column purified using QIAquick PCR Purification Kit (QIAGEN, Cat#28104). 3 μg of the purified PCR products was digested using 5 U of AarI (Thermo Fisher Scientific, Cat#ER1582) for 5 h at 37°C (***SI Appendix,* Table S3**), 5 U of Antarctic Phosphatase (New England Biolabs, Cat#M0289) for 1 h at 37°C (***SI Appendix,* Table S4**), and column purified. After purification, the digested PCR products were ligated into an AarI digested pRS316 pGPD ymScarlet expression vector using 3,200 U of T4 DNA Ligase (New England Biolabs, Cat#M0202) for 1 h at 25°C (***SI Appendix,* Table S5**). The ligation products were then transformed into NEB 5-alpha Competent E. coli (High Efficiency) (New England Biolabs, Cat#C2987) using 160 μl of ligation product in 1.6 ml of competent cells (32 x 50 μL reactions). Once transformed, the cells were pooled, inoculated into 500 ml of LB broth (Lennox) (Bioshop Canada, Cat#LBL405) in a 2 l flask containing 100 μg ml^-1^ of Carbenicillin (Wisent, Cat#400-112 IG), grown for 18 h, and maxiprepped using PureLink HiPure Plasmid Maxiprep Kit (Thermo Fisher Scientific, Cat#K210006). Test plating on LB agar (Lennox) plates with 100 μg ml^-1^ of Carbenicillin immediately after transformation estimated a total transformation yield of ∼1.9 x 10^5^ transformants. Sanger sequencing data from 93 randomly sampled colonies found a codon substitution rate of 1.065 ± 0.744 (***SI Appendix,* Datasets S1**).

### Pool Transformation of the Library into Yeast

The high-efficiency yeast transformation method used here was adapted from established protocols (109, 110) with modifications. Yeast strain JL701 was inoculated into 100 ml of YPD and incubated overnight for 18 h in a shaking incubator set to 30°C and 225 rpm. After 18 h, cells were refreshed to an OD_600_ of 0.4 in 400 ml of YPD and transferred back to the shaking incubator to be cultured for 4.5 h. Once refreshed, cells were pelleted by centrifugation, washed twice using 400 ml of 4°C water, once using 400 ml of 4°C electroporation buffer (1 M Sorbitol and 1 mM CaCl_2_), and resuspended in 80 ml of 25°C conditioning buffer (0.1 M Lithium acetate and 1 mM Dithiothreitol (DTT)). Resuspended cells were conditioned for 20 min in a shaking incubator set to 30°C and 225 rpm. After conditioning, cells were pelleted by centrifugation, washed once using 50 ml of 4°C electroporation buffer, and concentrated to a volume of 4 ml using 4°C electroporation buffer on ice. 50 ml of boiled 10 mg ml^-1^ UltraPure Salmon Sperm DNA Solution (Thermo Fisher Scientific, Cat#15632011) and 148 μg of the purified plasmid library was added to the resuspended cells and mixed well by pipetting. The mixture of cells and DNA was transferred into ice-chilled electroporation cuvettes with 2 mm gaps (Thermo Fisher Scientific, Cat#FB102) and electroporated at 2.5 kV, 25 mF, and 200 Ohms using a Gene Pulser II Electroporation System (Bio-Rad, Cat#165-2105). Immediately after electroporation, 1 ml of 4°C recovery media (1:1 ratio of 1 M Sorbitol and 2x YPD) was added directly into the electroporation cuvettes. Cells were then pooled and transferred to 80 ml of 4°C recovery media and incubated for 1 h in a shaking incubator set to 30°C and 150 rpm. After recovery, cells were pelleted by centrifugation, washed twice using 600 ml of water, and resuspended in 1 l of selective media comprising SD media minus Uracil plus 2% Glucose with 100 mg ml^-1^ of Carbenicillin (2 x 500 ml in 2 l flasks). A small aliquot of the transformed yeast was serially diluted and test-plated on SD media minus Uracil plus 2% Glucose agar plates immediately after transformation, estimating a total transformation yield of ∼3.6 x 10^7^ transformants. The remaining cells were grown for 30 h post-transformation in a shaking incubator set to 30°C and 225 rpm. This 30 h of outgrowth was crucial for two reasons. First, it provided sufficient time for variants to express, maturate, and for steady-state receptor abundance to stabilize (***SI Appendix* Fig. S11)**. Second, it provided sufficient time for the number of plasmids per cell to dilute to one **(*SI Appendix* Fig. S12**) (111).

### Fluorescence-Activated Cell Sorting

After 30 h of selective growth, yeast cells transformed with the site-saturation mutagenesis library were refreshed to an OD_600_ of 0.1, treated with 9-*cis-*retinal, and light/dark activated. Because refreshing the cells constituted a major population bottleneck, ∼24,800,000 cells were refreshed to preserve library diversity. For all FACS experiments, no Cyclohexamide was used to freeze protein synthesis in order to preserve cell viability. After light/dark activation, cells were pelleted, resuspended in selective media, and passed through 35 μm filter caps into 5 ml Falcon tubes (Corning) in a dark room. The cells were then transferred into 15 ml LiteSafe Light Blocking Centrifuge Tubes (Cole-Parmer, Cat#06344-59) and kept on ice in preparation for FACS. FACS was performed using a BD FACSymphony S6 SE (BD Biosciences) and the BD FACSDiva software (BD Biosciences, v9.8) using 488 and 561 nm lasers. Cells were sorted based on eGFP fluorescence into 3 bins to assess variant activity under dark conditions, eGFP fluorescence into 4 bins to assess variant activity under light conditions, and ymScarlet fluorescence into 5 bins to assess per cell receptor abundance. To minimize uncontrolled rhodopsin variant activation due to ambient light during FACS, especially for the dark treatment condition, cells were split into multiple LiteSafe Light Blocking Centrifuge Tubes (Cole-Parmer, Cat#06344-59), temperature controlled at 4°C, and never sorted for more than 15 min after removing centrifuge tube lids outside the dark room. After FACS, cells were pelleted by centrifugation and resuspended in selective media at a concentration of 100,000 cells ml^-1^ and grown overnight for 24 h in a shaking incubator set to 30°C and 225 rpm. Test plating of sorted cells immediately after FACS on SD media minus Uracil plus 2% Glucose agar plates estimated that ∼25-30 million cells were collected for each cell sorting experiment. Example gating strategy is provided (***SI Appendix* Fig. S13**).

### Plasmid Extraction from Sorted Cells

After 24 h of selective growth, sorted cells were pelleted by centrifugation and resuspended in 1.2 ml of yeast plasmid extraction buffer (1 M Sorbitol (Bioshop Canada, Cat#SOR508), 100 mM EDTA (Millipore Sigma, Cat#324503), 14 mM DTT (Thermo Fisher Scientific, Cat#R0861), and 1,000 U of 2-Mercaptoethanol-activated Zymolyase (Bioshop Canada, Cat#ZYM001). The resuspensions were incubated for 2 h in a shaking incubator set to 30°C and 100 rpm to digest yeast cell walls and form spheroplasts. Plasmids were extracted using Zymoprep Yeast Plasmid Miniprep II (Zymo Research, Cat#D2004), with the reagent volumes scaled up 5x, and eluted in 20 μl of 37°C Tris-EDTA (pH 8) buffer (Thermo Fisher Scientific, Cat#327345000).

### Amplicon Library Preparation

Plasmids extracted from sorted yeast cells were used as the DNA template for generating amplicon libraries for deep sequencing. A set of 10 PAGE Ultramer DNA oligonucleotides, synthesized by Integrated DNA Technologies, were designed to amplify the sequence in *RHO* encoding EL2–TM6 and to add all the necessary elements for direct sequencing on Illumina flow cells (***SI Appendix,* Table S6**). Each oligonucleotide in the set contains an Illumina adapter sequence, a unique index barcode, a sequencing primer binding sequence, heterogeneity spacer of variable length, and an insert binding sequence. The heterogeneity spacers were designed using a custom script (112) optimized to counteract the issue of low nucleotide diversity during Illumina sequencing. The oligonucleotides were arranged in a 5 x 5 Illumina combinatorial index matrix to enable multiplexed sequencing of 25 samples, which were populated by plasmids extracted from each of the cell sorting bins. Amplicons were generated with 18 cycles of PCR using NEBNext Ultra II Q5 Master Mix (New England Biolabs, Cat#M0544) (***SI Appendix,* Table S7**).

### Deep Sequencing

Amplicon libraries were processed and deep sequenced by the Centre for the Analysis of Genome Evolution and Function at the University of Toronto. Amplicons were purified using 0.8x ratio of MGIeasy DNA Clean Beads (Complete Genomics, Cat#1000005278) and quantified using Qubit 2.0 Fluorometer (Life Technologies, Cat#Q32866) with Qubit dsDNA HS Kit (Thermo Fisher Scientific, Cat#Q32851). Samples were pooled in equal amounts, then denatured and diluted to a sequencing concentration of 750 pM. Sequencing was performed on an Illumina NextSeq 2000 Sequencing System (Illumina, Cat#20038897) using the P2 chemistry with 300 bp Paired-End reads (Illumina, Cat#20075295). The raw data was converted to FASTQ format using BCL convert (Illumina, v4.2.7). Five percent PhiX control (Illumina, Cat#FC-110-3001) was included during sequencing.

### Deep Sequencing Data Processing

Deep sequencing reads were processed using a custom computational workflow on an AMD Ryzen 5950X CPU (Advanced Micro Devices, Product ID: 100-100000059WOF) equipped with a Nvidia GeForce RTX 3090 GPU (Nvidia, Product ID: 900-1G136-2510-000). Reads were trimmed using Cutadapt (v4.4) (113) to remove primer sequences at the beginning and end of reads, leaving only the region that encodes EL2–TM6. Trimmed reads were merged using NGmerge (v0.3) (114) and quality filtered using fastp (v.0.24.0) (115) and SeqKit (v2.8.2) (116) to remove reads of low quality (per read mean Phred score < 33), unexpected length, or contain ambiguous base calls (***SI Appendix,* Fig. S5*C***). The remaining reads were downsampled to match the frequencies of cells sorted into each FACS bin (***SI Appendix,* Fig. S5*D***). From the downsampled reads, variants were called, recorded in variant calling format, and filtered based on abundance to remove variants that were called < 15x. The remaining variant calls were then processed to generate CSV files containing binned distributions (the proportion of each variant across FACS bins) and mixing coefficients (the proportion of each variant in the library). These files, together with CSVs containing gating boundaries specific to each cell sorting experiment and the overall fluorescence intensity distributions of the library acquired during FACS, were inputted into dSort-Seq (117) to calculate variant effect scores. A bootstrap-analysis was performed, from the read downsampling step to calculating variant effect scores for the purpose of (i) reducing potential bottlenecks introduced during read downsampling and (ii) accounting for stochasticity in the two-component log-Gaussian mixture model implemented in dSort-Seq. The bootstrap-analysis was repeated 30x. Variant effect scores calculated from each analysis were averaged across bootstraps and scaled such that nonsense and synonymous variants were scaled to 0 and 1, respectively, to generate final variant effect scores (***SI Appendix,* Fig. S5*E*-*F***). Metadata of calculated variant effect scores is provided (***SI Appendix,* Dataset S2**).

### Statistics and Reproducibility

All graphing and statistical analyses were performed in GraphPad Prism (v10.5.0), FlowJo (v10.10.0), or Microsoft Excel (Microsoft Office v16.97.2). *P* < 0.05 was considered statistically significant.

## Data, Materials, and Software Availability

Deep sequencing datasets have been deposited in the European Nucleotide Archive with the BioProject accession number PRJEB100940. All original code used to process and analyze the deep sequencing data have been deposited on GitHub and is publicly available at https://github.com/jingliu-ut/SynLib_NextSeq_DMS_Analysis.

## Acknowledgments

We thank all members of the Chang Lab for helpful discussions. We are grateful to Frederick P. Roth of the University of Toronto Department of Molecular Genetics and the Lunenfeld-Tanenbaum Research Institute (now at the University of Pittsburgh Department of Computational and Systems Biology) and Jane Mitchell of the University of Toronto Department of Pharmacology and Toxicology for comments and suggestions; Nathalie Simard and Parva Thakker of the University of Toronto Temerty Faculty of Medicine Flow Cytometry Facility for assistance with flow cytometry and cell sorting; Sergey V. Plotnikov, Ernest Iu, and Fernando R. Valencia of the University of Toronto Department of Cell and Systems Biology for providing expertise and access to fluorescence microscopes; John A. Calarco of the University of Toronto Department of Cell and Systems Biology for access to an electroporation system; David S. Guttman, Pauline Wang, and Lijie Yuan of the University of Toronto Centre for the Analysis of Genome Evolution and Function for deep sequencing advice, strategies, and support.

This work was supported by a NSERC Discovery grant to B.S.W.C., an Ontario Graduate Scholarship and a Vision Science Research Program Fellowship to S.K.C., and a Vision Science Research Program Fellowship to J.L.; Figs. 1A,C were created in part using biorender.com.

## Supplemental Information

**Figure S1.**
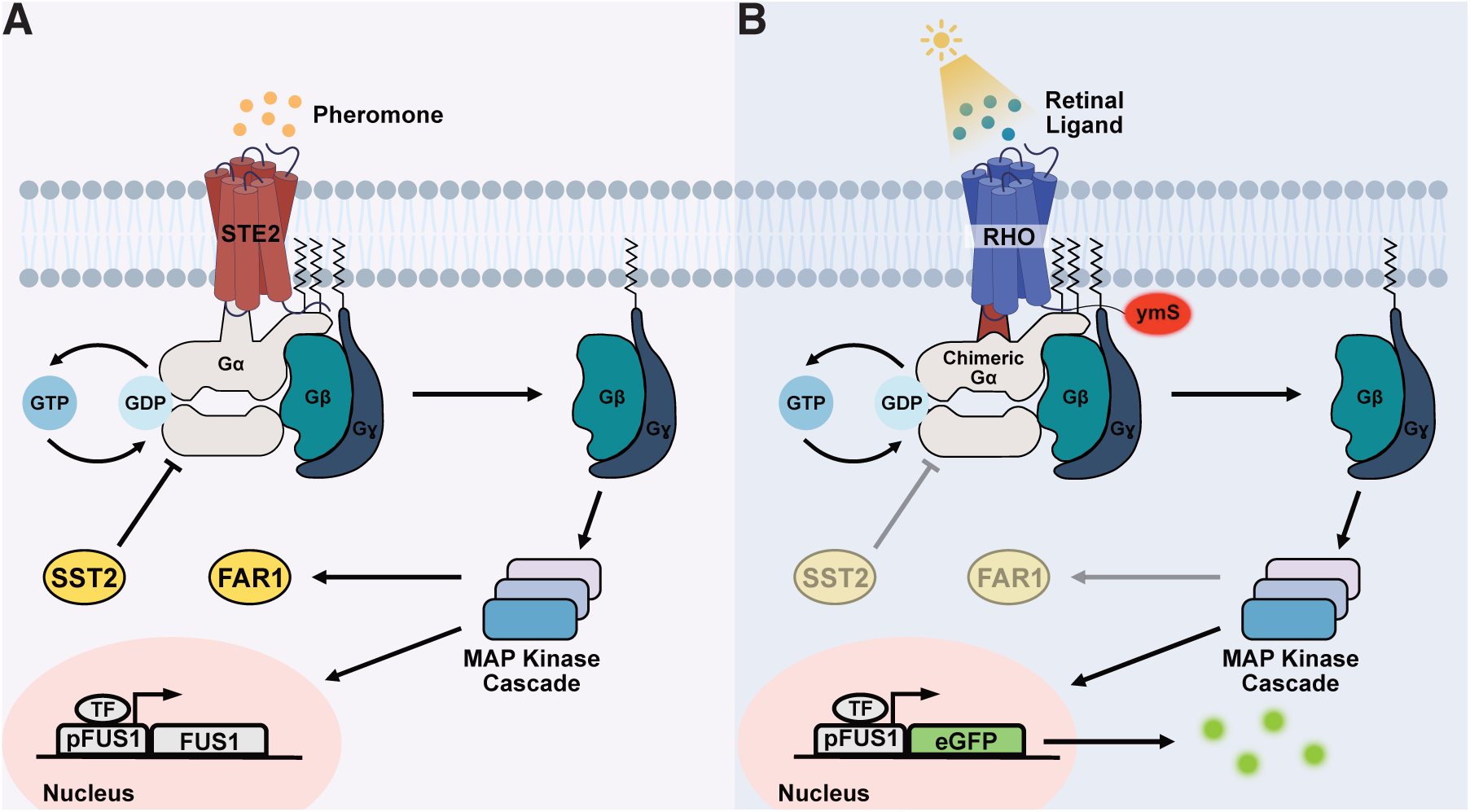
Engineered GPCR signaling pathway in yeast enables simultaneous tracking of rhodopsin variant activation and abundance. **(A)** The mating response pathway endogenous to *S. cerevisiae*. The yeast GPCR STE2 is activated by pheromone produced by yeast of the same species but opposite mating type. Upon pheromone binding, STE2 adopts its active conformation, triggering GDP-GTP exchange at the Gα G protein and release of the Gβγ dimer. Once released, Gβγ activates MAP kinase signaling, leading to upregulation of mating genes. **(B)** A modified *S. cerevisiae* mating pathway engineered to functionally couple heterologously expressed mammalian GPCRs, enabling their activity to be measured via a pathway-responsive fluorescent reporter. Key modifications include (i) replacing STE2 with a heterologously expressed mammalian GPCR (here, human rhodopsin (RHO)); (ii) replacing the endogenous Gα subunit with a chimeric Gα that couples the pathway to rhodopsin; (iii) removing SST2, a negative regulator of G protein signaling; (iv) removing FAR1, an inducer of cell-cycle arrest upon pathway activation; (v) integrating a pathway-responsive eGFP fluorescent reporter. Fusion of rhodopsin to the fluorescent protein yeast codon-optimized mScarlet (ymS) enables monitoring of cellular rhodopsin abundance.

**Figure S2.**
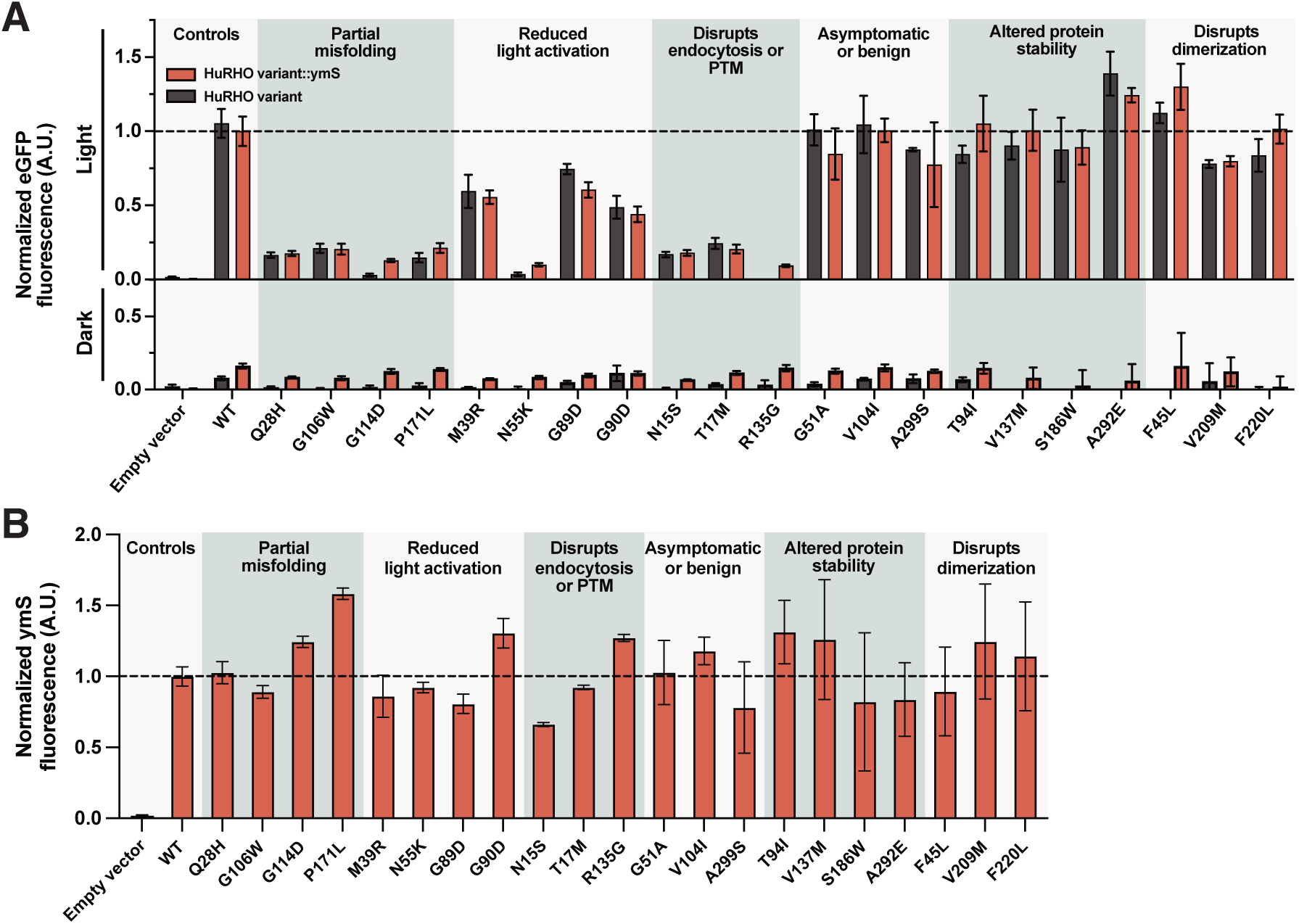
Activity and cellular abundance measurements of 21 rhodopsin variants in yeast. **(A)** Rhodopsin-mediated signal transduction, measured as eGFP fluorescence, under activating (light) and non-activating (dark) conditions. Missense variants are classified according to experimentally determined biochemical or cellular characteristics, including variants that cause partial misfolding, reduce activation, disrupt endocytosis or post-translational modifications (PTM), are asymptomatic, alter protein stability, or disrupt receptor dimerization. For each variant signal transduction is shown for ymScarlet-fused (HuRHO variant-ymS, red bars) and non-fused (HuRHO variant, black bars) versions. Bars represent mean ± standard deviation of n = 4 biological replicates with each replicate comprising data from 10,000 cells acquired using flow cytometry. **(B)** Cellular abundance for each variant in (A), measured as ymScarlet (ymS) fluorescence. Bars represent mean ± standard deviation of n = 4 biological replicates with each replicate comprising data from 10,000 cells acquired using flow cytometry.

**Figure S3.**
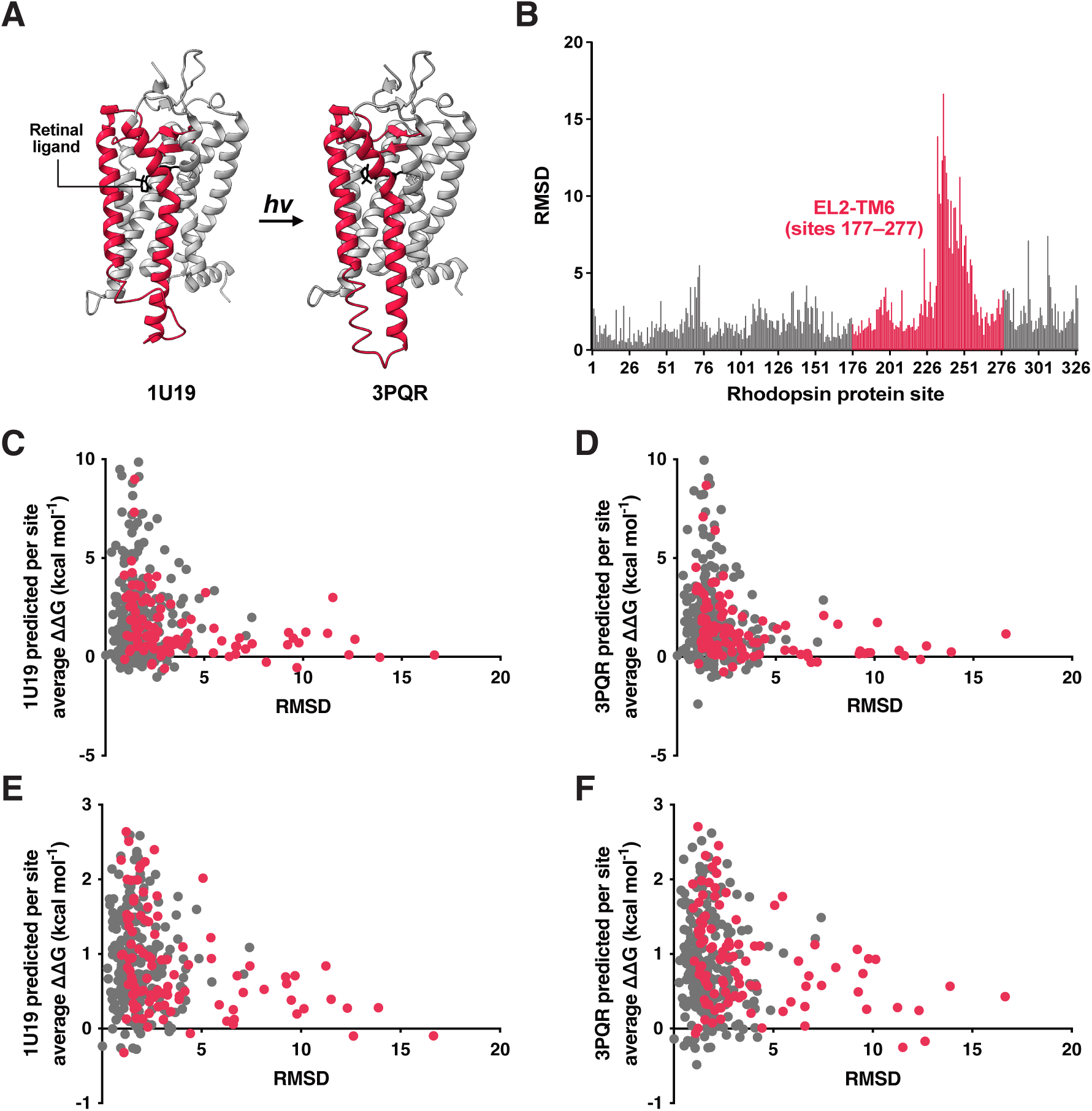
Site-specific comparison of predicted mutational effects on rhodopsin stability and structural mobility during activation. **(A)** Inactive/dark-state (PDB: 1U19) and active-state (PDB: 3PQR) rhodopsin structures. Extracellular loop 2 (EL2), helix 5 (H5), intracellular loop 3 (ICL3), and helix 6 (H6) are coloured in crimson. **(B)** Per residue Cα root-mean-squared deviation (RMSD) between inactive- and active-state structures, calculated using UCSF ChimeraX (1). **(C-D)** Per-site RMSD (from B) plotted against the mean predicted ΔΔG across all possible amino acid substitutions at each site, calculated using FoldX (2) using the inactive-state (C) and active-state (D) structures. **(E-F)** As in (C-D), using PoPMuSiC (3) in place of FoldX to predict ΔΔG.

**Figure S4.**
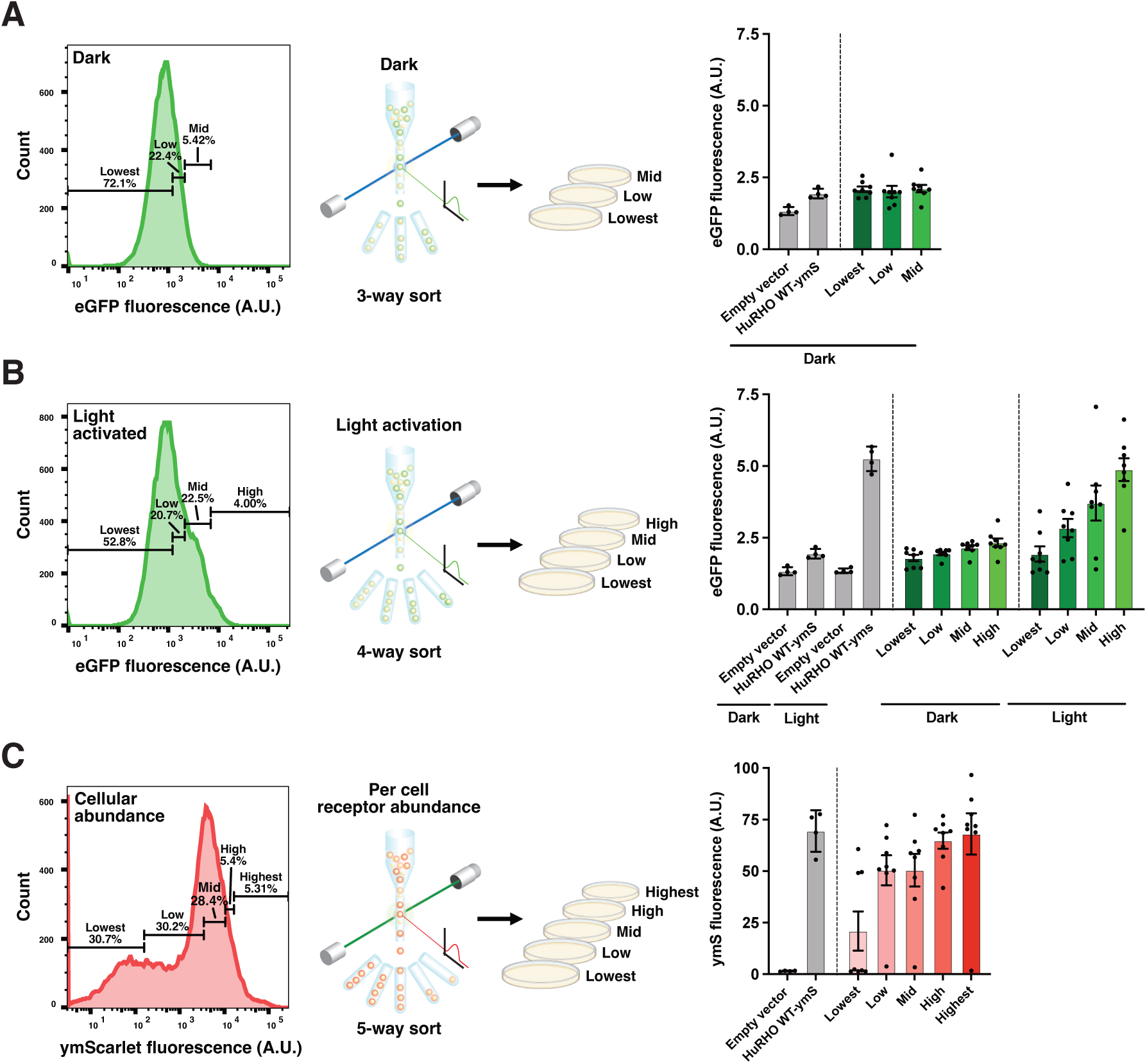
Rhodopsin variant activity and cellular abundance across FACS bins. **(A)** Dark-treated cells were sorted by eGFP fluorescence into three bins (Lowest, Low, Mid), with the percentage of cells sorting into each bin indicated. A small number of cells from each bin were plated on selective agar, and eight colonies were randomly picked per plate and assayed for dark-state rhodopsin activity, measured as eGFP fluorescence. Bar graph data represent mean ± standard deviation for empty vector and wild-type human rhodopsin (HuRHO WT-ymS) controls (n = 4 biological replicates) and randomly picked colonies (n = 8 per agar plate). Each dot represents data from 10,000 cells from a single colony. **(B)** As in (A), except cells were light-activated and sorted into four bins (Lowest, Low, Mid, High), and randomly picked colonies were assayed under both dark and light conditions. **(C)** As in (A), except cells were sorted by cellular receptor abundance, measured as ymScarlet (ymS) fluorescence, into five bins (Lowest, Low, Mid, High, Highest).

**Figure S5.**
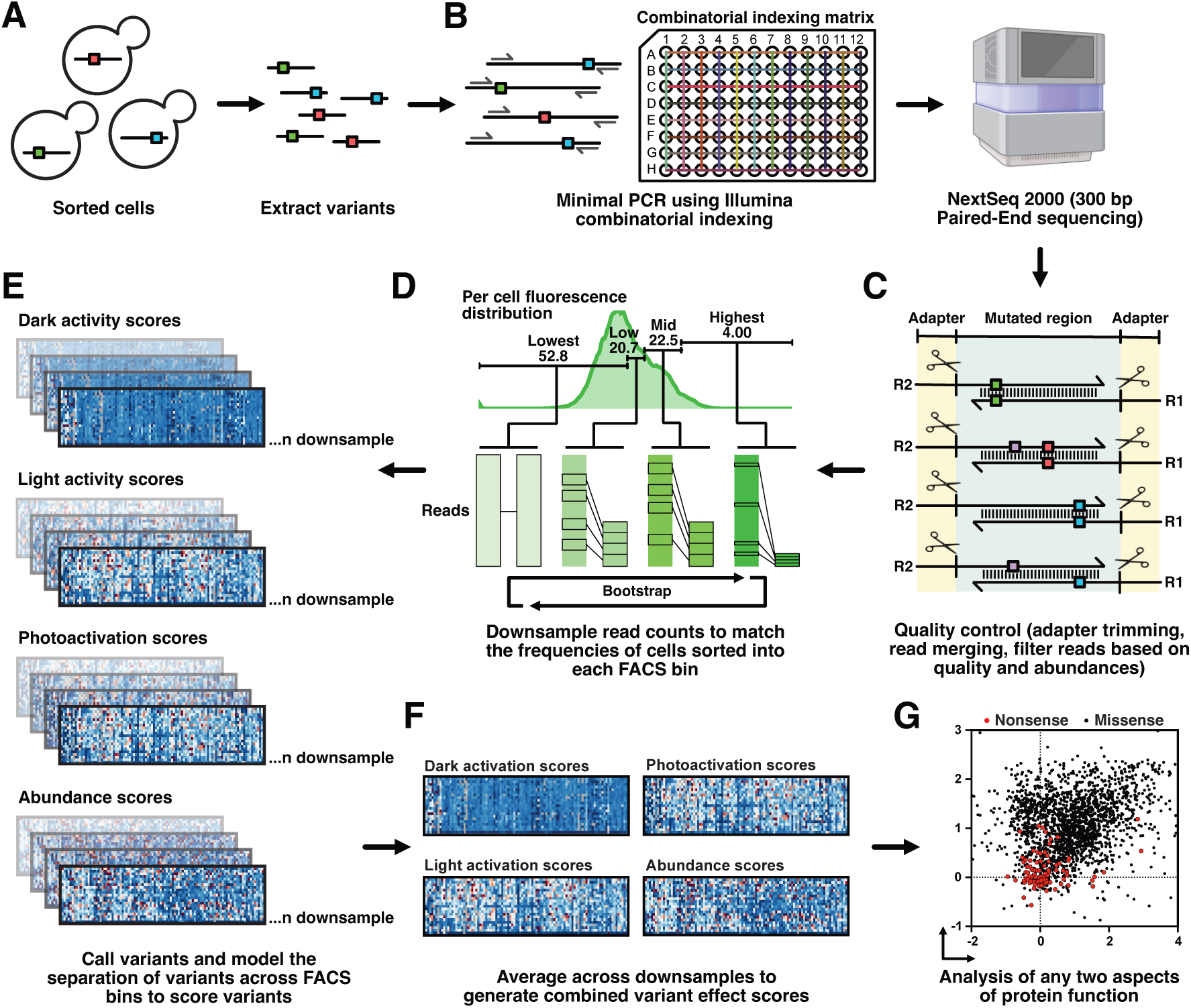
FACS-Seq post-cell sorting workflow. **(A)** RHO variants were extracted from FACS-sorted yeast cells. **(B)** Extracted variants were amplified using minimal-cycle PCR to add the elements required for direct sequencing on Illumina flow cells. These include AmpliSeq adapters, sequencing primer binding regions, index barcodes, and heterogeneity spacers. Amplicson were sequenced on an Illumina NextSeq 2000 using 300 bp Paired-End sequencing. **(C)** Reads were processed through quality-control steps including adapter trimming, Paired-End read merging, and filtering based on read quality and variant abundance. **(D)** Merged reads from each FACS bin were downsampled to match the frequency of cells sorted into each bin. For each FACS experiment, downsampling was performed 30 times to account for downsampling bias and bottleneck effects. **(E)** Variants were called from the downsampled reads and modeled based on their separations across FACS bins to generate variant effect scores. This process was performed independently for each downsample. **(F)** Variant effects scores from individual downsample analyses were averaged to generate combined variant effect scores and final variant effect maps. **(G)** Downstream analysis may include comparing pairs of protein properties.

**Figure S6.**
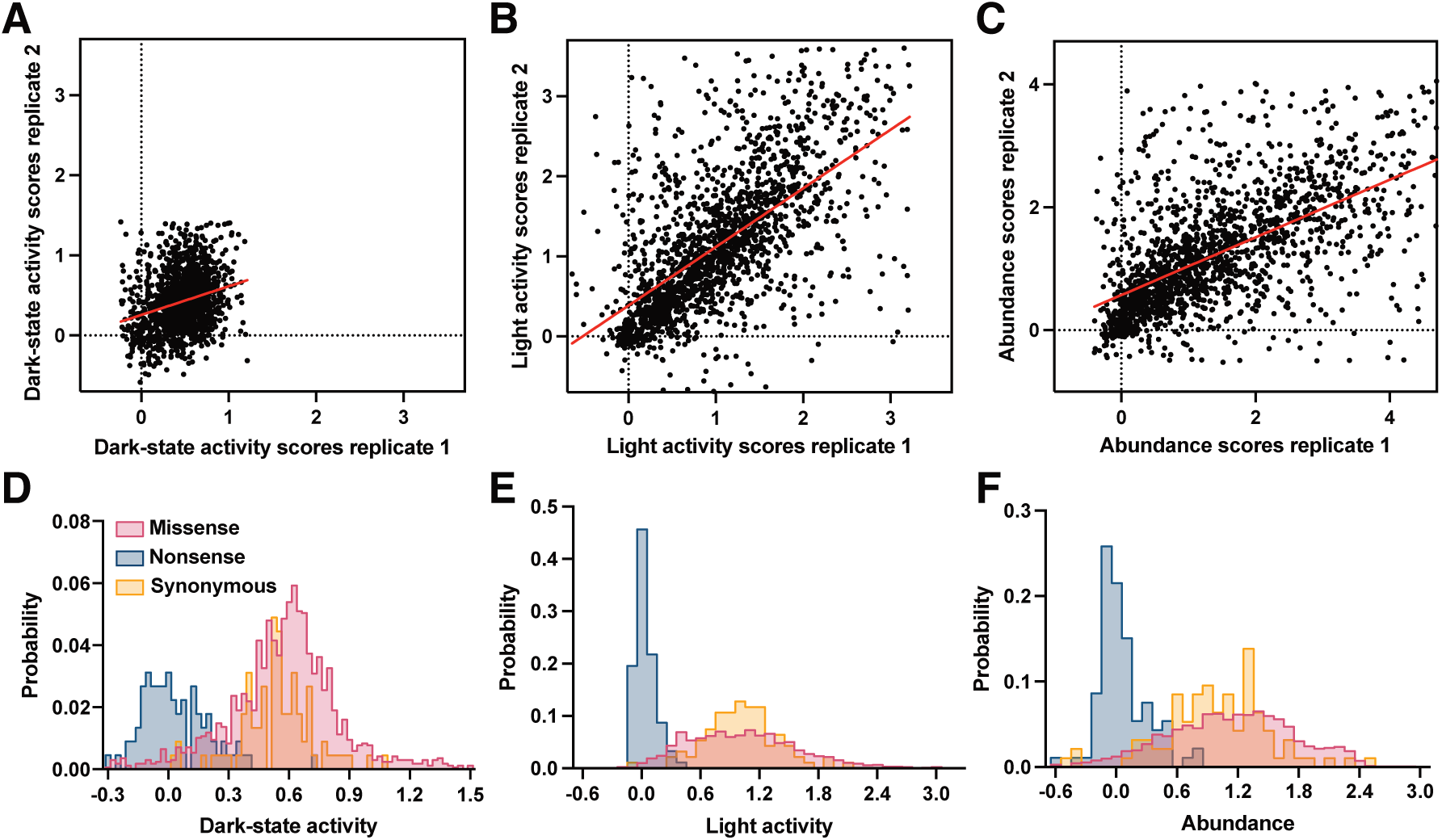
DMS replication analysis and distributions of variant effects. **(A-C)** Scatter plots showing variant effect scores from two biological replicate experiments for variant activity in the dark (Pearson’s r = 0.2605, Spearman’s r = 0.2625) (A), variant activity when exposed to light (Pearson’s r = 0.6107, Spearman’s r = 0.6474) (B), and variant abundance (Pearson’s r = 0.5895, Spearman’s r = 0.6282) (C). **(D-F)** Distributions of variant effects for variant activity in the dark (D), variant activity when exposed to light (E), and variant abundance (F). In all cases, distributions between synonymous and missense variants are distinct (two-sided Mann-Whitney U test P < 0.0001).

**Figure S7.**
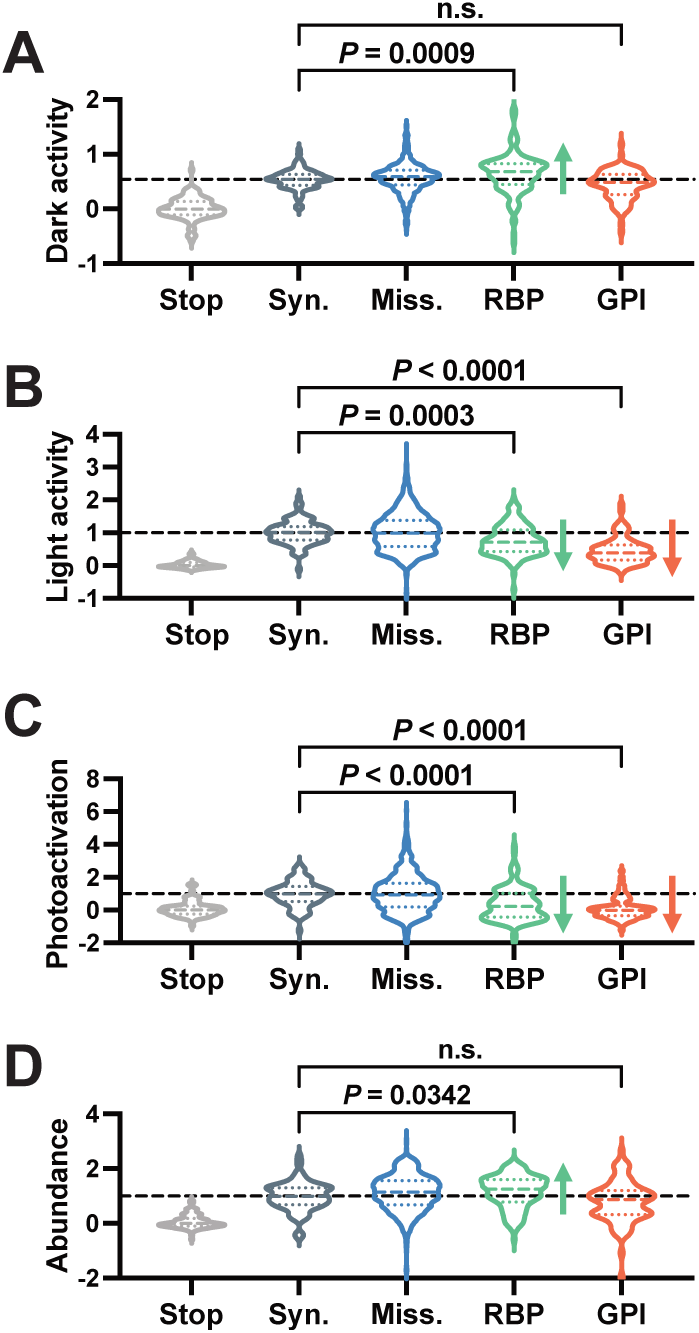
Amino acid substitutions in the RBP and GPI show asymmetric pleiotropic effects. **(A)** Dark-state activity is significantly elevated for amino acid substitutions of RBP residues (*P* = 0.0009), but not for GPI residues. **(B)** Activity when exposed to light is significantly reduced for both RBP (*P* = 0.0003) and GPI residues (*P* < 0.0001). **(C)** Photoactivation is significantly reduced for both substitutions of RBP and GPI residues (*P* < 0.0001). **(D)** Abundance is significantly elevated for substitutions of RBP residues (*P* = 0.0342), but not for GPI residues. Stop denotes premature stop codons, Syn. denotes synonymous variants, and Miss. denotes all missense variants. *P* values were determined using Kruskal-Wallis test with Benjamini-Hochberg correction. n.s. denotes not significant (*P* > 0.05).

**Figure S8.**
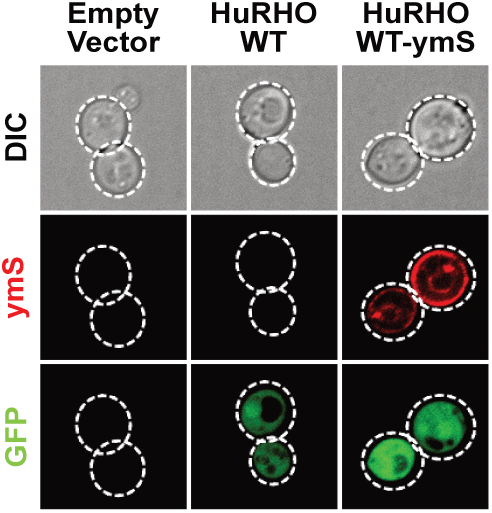
Representative differential interference contrast (DIC) and fluorescent images of rhodopsin localization (ymS) and rhodopsin-mediated pathway activation (eGFP) in yeast. Images were acquired after treating the cells with 9*-cis-*retinal and light. These fluorescence images show that rhodopsin is localized to the yeast plasma membrane and activates the pathway-responsive eGFP fluorescence reporter in response to treatment with retinal and light.

**Figure S9.**
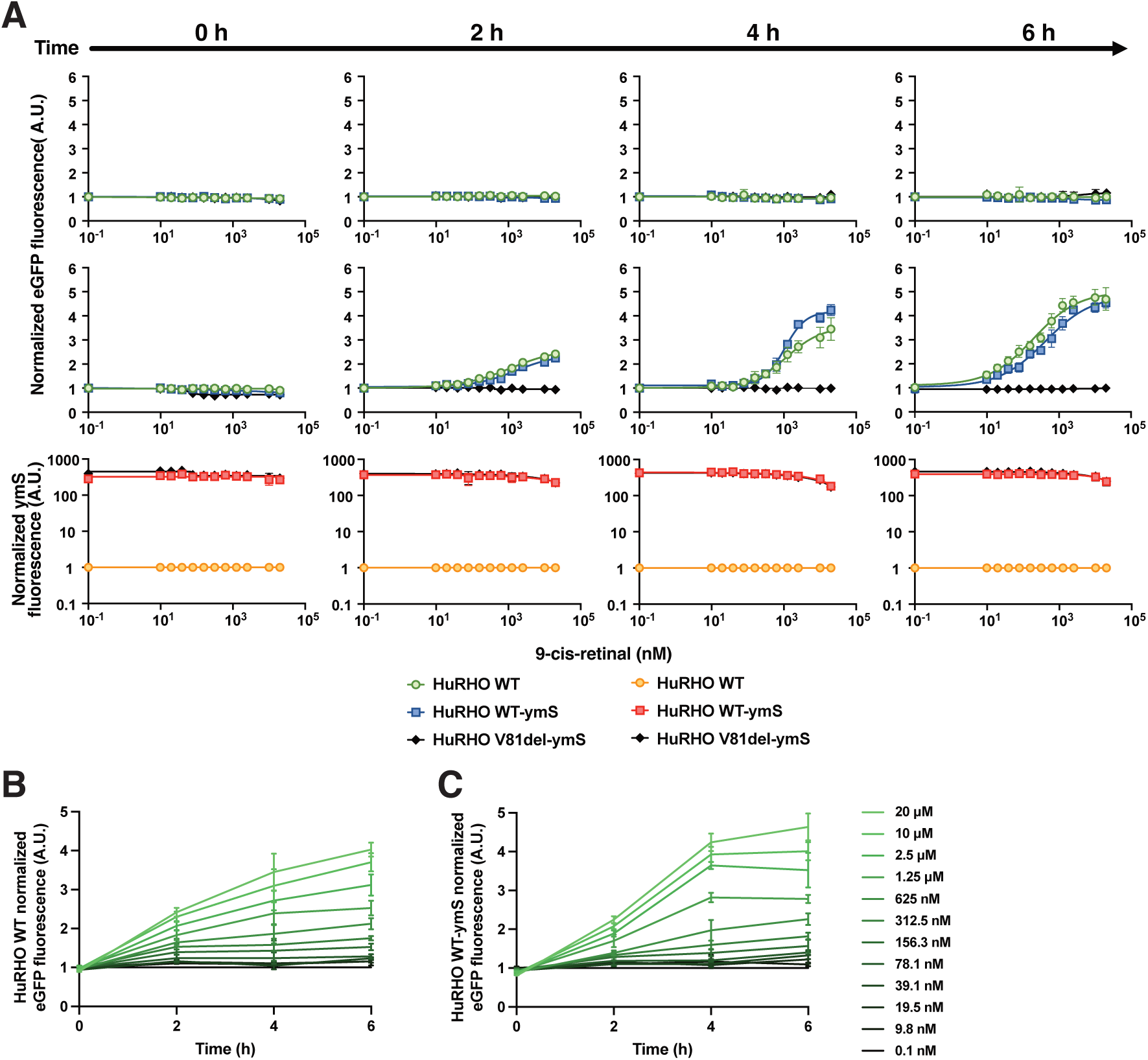
Time-resolved dose-response of rhodopsin photoactivation and cellular abundance in response to retinal and light. **(A)** Dose-response curves showing rhodopsin activation and cellular abundance as a function of 9*-cis-*retinal concentration over time. Yeast strains constitutivively expressing wild-type human rhodopsin (HuRHO WT), ymScarlet-fused wild-type human rhodopsin (HuRHO WT-ymS), or the loss-of-function variant HuRHO V81del-ymS were treated with 9*-cis-*retinal (0.1 nM to 20 μM) and kept in the dark (top row) or exposed to light (middle and bottom rows). Rhodopsin activation (eGFP) fluorescence and cellular rhodopsin abundance (ymScarlet fluorescence) were measured every 2 h for each condition. Each point represents mean ± standard deviation of n = 4 biological replicates, each comprising 10,000 cells acquired by flow cytometry. **(B-C)** Analysis of the data from (A) to show changes in rhodopsin activity over time and across 9*-cis-*retinal concentrations.

**Figure S10.**
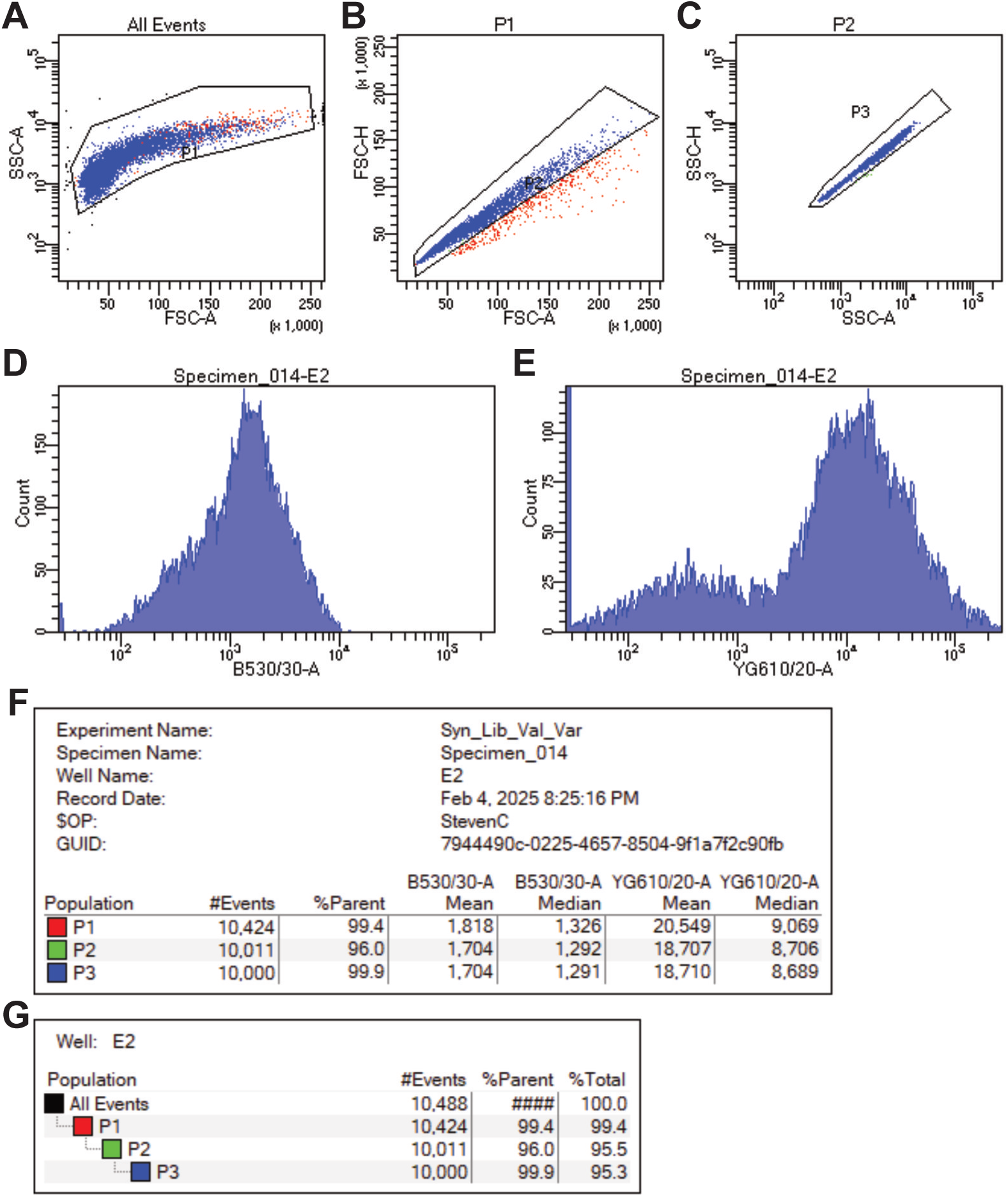
Example gating hierarchy for isolating singlets. **(A)** Gating for yeast cells using Side Scatter Area (SSC-A) and Forward Scatter Area (FSC-A). **(B-C)** Gating for singlets using Forward Scatter Height (FSC-H) and Forward Scatter Area (FSC-A) (B) together with Side Scatter Area (SSC-H) and Side Scatter Height (SSC-A) (C). **(D)** Histograms showing the distribution of receptor activation as a function of eGFP fluorescence acquired using the Blue 488 nm laser with a 530/30 detection filter (B530/30-A). **(E)** Histograms showing the distribution of receptor activation as a function of eGFP fluorescence acquired using the Yellow-Green 561 nm laser with a 610/20 detection filter (YG610/20-A). **(F)** Live gating statistics. **(G)** Gating hierarchy.

**Figure S11.**
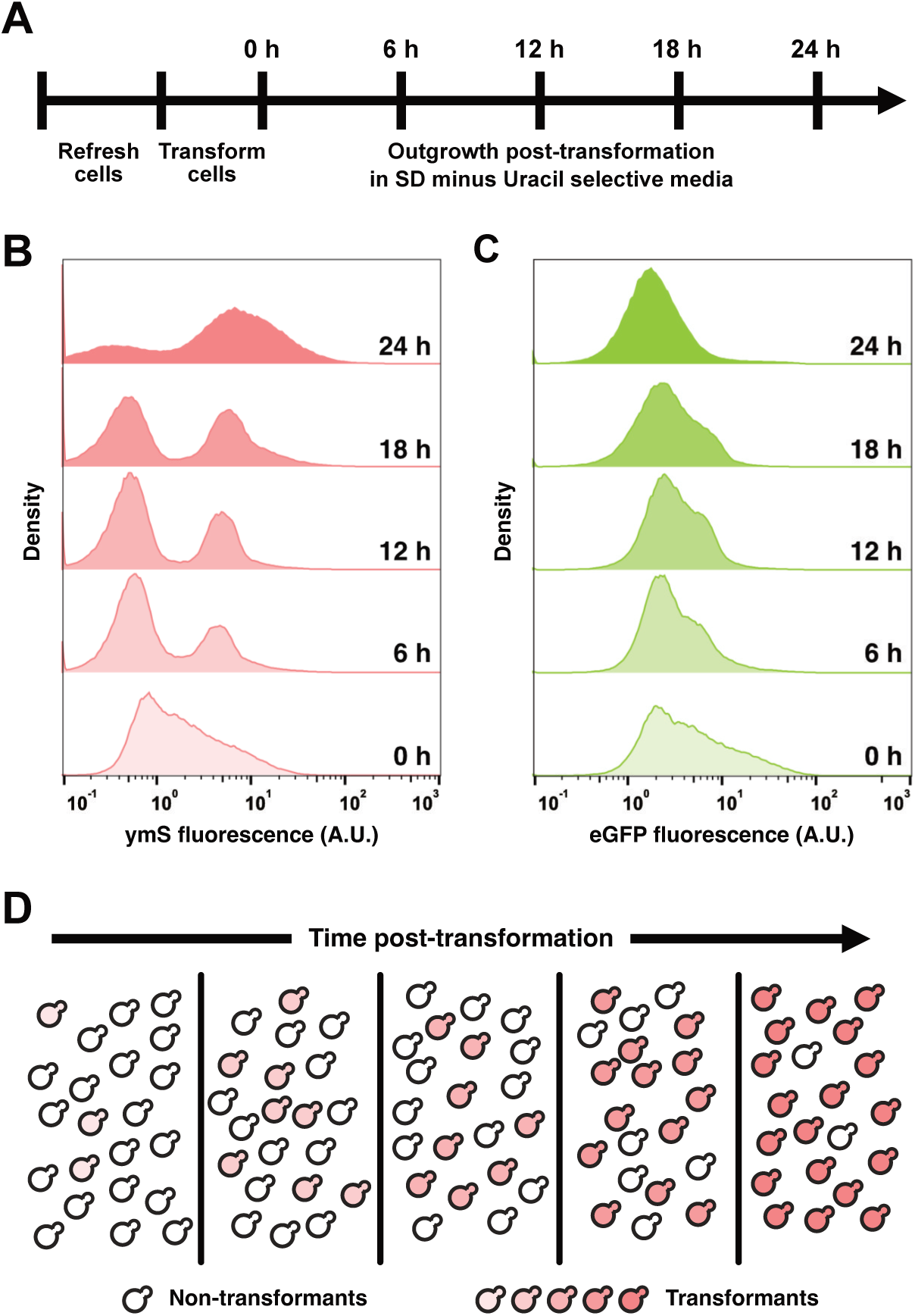
Transformed yeast cells expand to dominate the population culture over 24 hours of selective growth. **(A)** Timeline showing transformation of yeast cells with a plasmid library for expressing human rhodopsin variants followed by outgrowth in selective media. Each plasmid in the library contains a URA3 selection marker, which confers growth under selection in synthetic dextrose media deficient in Uracil, and a centromeric origin of replication for maintaining the number of plasmids per cell at or near one. **(B)** Distribution of cells expressing rhodopsin variants as a function of per cell ymScarlet (ymS) fluorescence over time. Since each rhodopsin variant in the plasmid library is fused to ymS, a right-shift in ymScarlet fluorescence indicates expansion of transformed yeast cells. Distribution data represent per cell fluorescence measurements of 100,000 cells acquired using flow cytometry at 0, 6, 12, 18, and 24 h timepoints post-transformation. **(C)** Distributions showing the per cell eGFP fluorescence of the cells in (B). **(D)** Schematic representation showing a subpopulation of transformants overtaking non-transformants under selective conditions over time.

**Figure S12.**
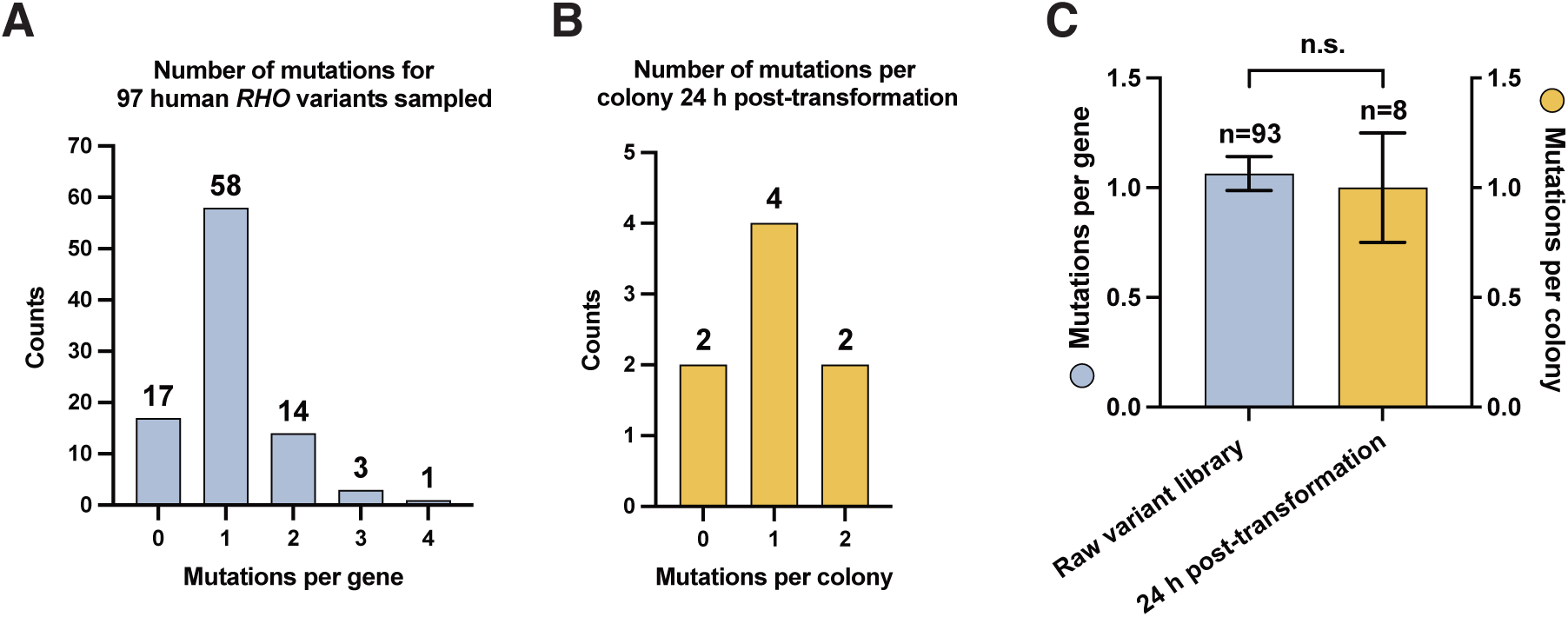
Controlling for multiple vector transformants. **(A)** Number of single-codon substitutions per gene for 93 randomly sampled RHO sequences from the site-saturation mutagenesis library covering EL2-H6. **(B)** Number of single-codon substitutions per gene of plasmids extracted from 8 randomly sampled yeast colonies. These colonies were from yeast cells that were transformed using the site-saturation mutagenesis library, grown for 24 h in the appropriate synthetic dextrose minus uracil selection media (SD-URA), and plated on SD-URA plates. **(C)** Comparison of the number of mutations per RHO gene in the library from (A) versus the number of mutations per RHO gene for each colony sampled from (B). Error bars represent standard deviations. Statistical significance was determined using a two-sided Student’s t-test. n.s. denotes not significant.

**Figure S13.**
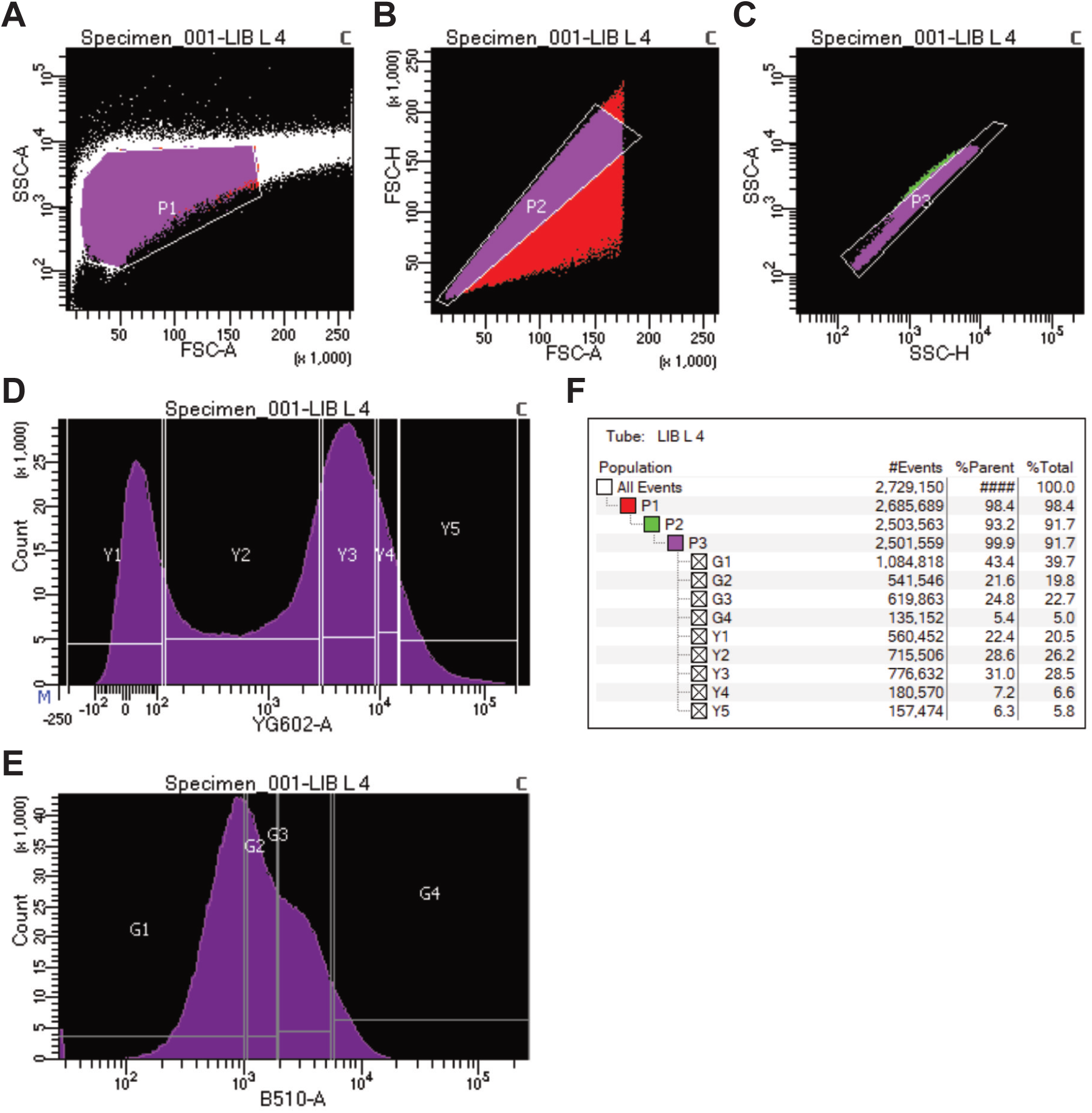
Example FACS gating strategy. **(A)** Gating for yeast cells using Side Scatter Area (SSC-A) and Forward Scatter Area (FSC-A). **(B-C)** Gating for singlets using Forward Scatter Height (FSC-H) and Forward Scatter Area (FSC-A) together with Side Scatter Area (SSC-A) and Side Scatter Height (SSC-H). **(D)** Gating based on ymScarlet fluorescence for per cell variant abundance using the Yellow-Green 561 nm laser with a 602/40 detection filter (YG602-A). **(E)** Gating based on eGFP fluorescence for variant activation using the Blue 488 nm laser with a 510/20 detection filter (B510-A). **(F)** Gating hierarchy.

**Table S1.**
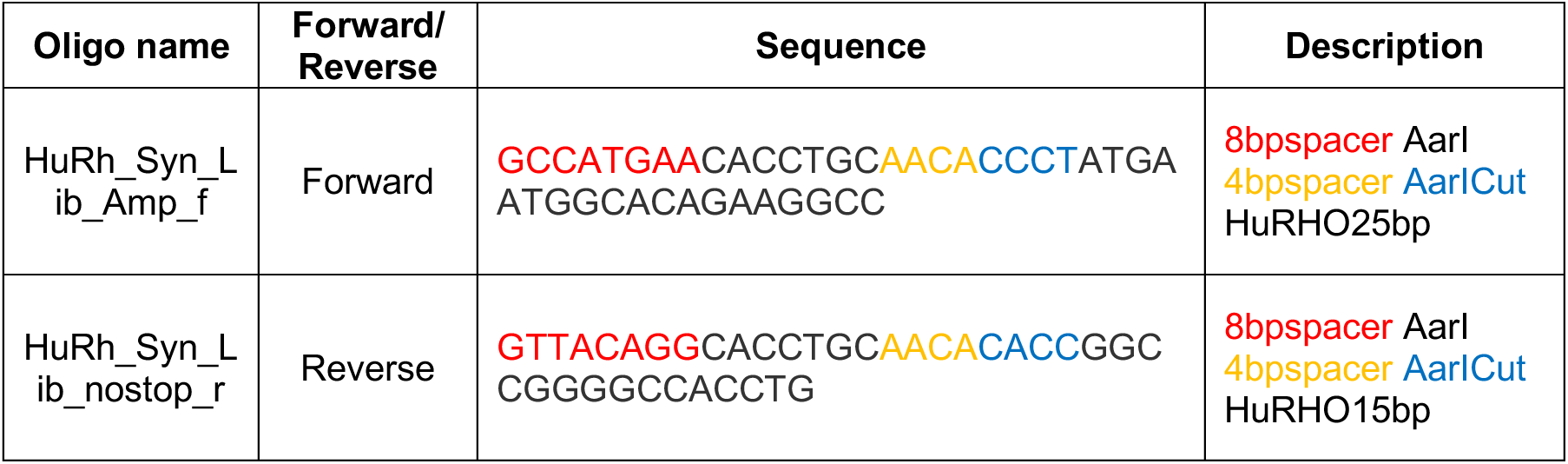
Oligonucleotides for cloning *RHO* variant library synthesized by Ranomics.

**Table S2.** PCR reagents and cycling conditions for adding flanking AarI cut sites to the *RHO* variant library synthesized by Ranomics.

| Kit: KOD Xtreme Hot Start DNA Polymerase (Millipore Sigma; 71975-M) |  |  |  |
| --- | --- | --- | --- |
|  | Name | Concentration | Volume (μL) |
| DNA | Ranomics_Syn_Lib | 10 ng/μL | 1 |
| Buffer | KOD Xtreme PCR Buffer | 2x | 25 |
| dNTPs | dNTPs | 2 mM | 10 |
| Oligo F | HuRh_Syn_Lib_Amp_f | 10 μM | 1.5 |
| Oligo R | HuRh_Syn_Lib_nostop_r | 10 μM | 1.5 |
| Enzyme | KOD Xtreme Hot Start DNA Polymerase | 1 U/μL | 1 |
| Water |  |  | 10 |
|  |  | **Total volume** | 50 |
| Time | Temperature (°C) | Number of cycles |  |
| 2 min | 94 | 1x |  |
| 10 sec | 98 | 25x |  |
| 30 sec | 70; decrease by 0.5 per cycle |  |  |
| 1 min 30 sec | 68 |  |  |
| 7 min | 68 | 1x |  |
| Hold | 4 |  |  |

**Table S3.**
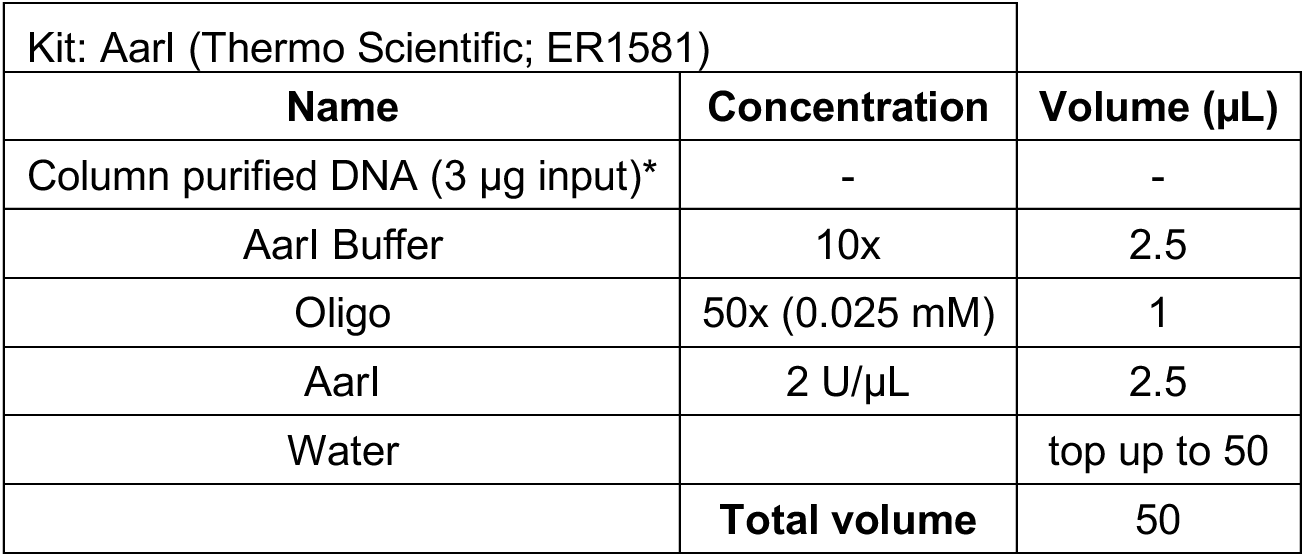
AarI digestion for cloning the *RHO* variant library into a centromeric yeast expression vector.

| Kit: AarI (Thermo Scientific; ER1581) |  |  |
| --- | --- | --- |
| Name | Concentration | Volume (μL) |
| Column purified DNA (3 μg input)* | - | - |
| AarI Buffer | 10x | 2.5 |
| Oligo | 50x (0.025 mM) | 1 |
| AarI | 2 U/μL | 2.5 |
| Water |  | top up to 50 |
|  | **Total volume** | 50 |

**Table S4.** Antarctic phosphatase digestion.

| Kit: Antarctic Phosphatase (New England Biolabs; M0289) |  |  |
| --- | --- | --- |
| Name | Concentration | Volume (μL) |
| AarI Digested DNA |  | 50 |
| Antarctic Phosphatase Buffer | 10x | 6 |
| Antarctic Phosphatase | 5 U/μL | 1 |
| Water |  | 3 |
|  | **Total volume** | 60 |

**Table S5.** T4 DNA ligase ligation reaction.

| Kit: T4 DNA Ligase (New England Biolabs; M0202) |  |  |
| --- | --- | --- |
| Name | Concentration | Volume (μL) |
| Column purified DNA insert (390.4 ng input)* | - | - |
| Column purified linear vector (800 ng input)* | - | - |
| 2x Ligase Buffer |  | 16 |
| T4 DNA Ligase | 400 U/μL | 8 |
| Water |  | top up to 160 |
|  | **Total volume** | 160 |

**Table S6.**
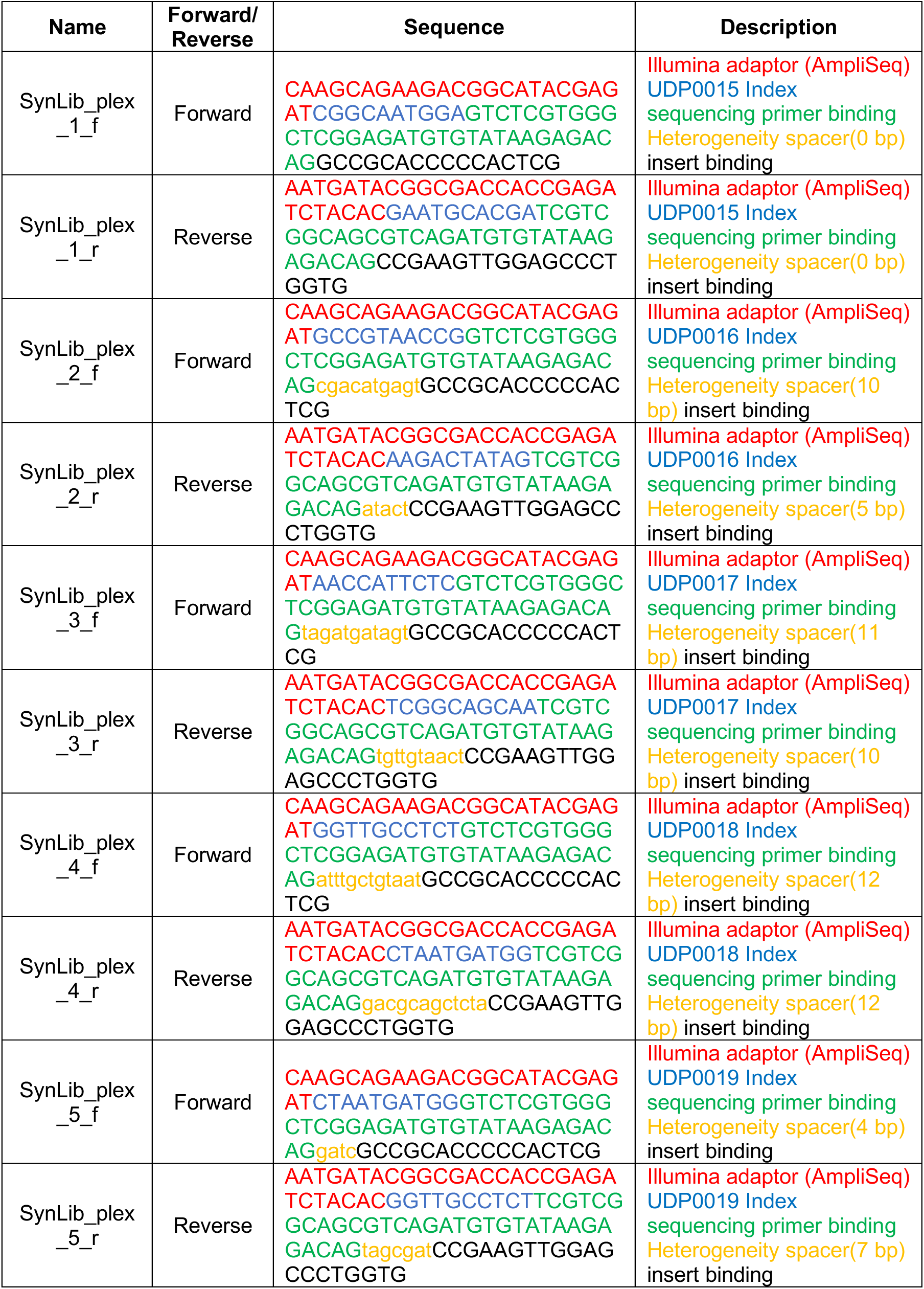
PAGE Ultramers for generating amplicons for deep sequencing.

**Table S7.** PCR reagents and cycling conditions for generating amplicons for deep sequencing.

|  |  |  |  |
| --- | --- | --- | --- |
| Kit: NEBNext Ultra II Q5 Master Mix (M0544) |  |  |  |
|  | **Name** | **Concentration** | **Volume (μL)** |
| DNA | Plasmid extracted from sorted cells | - | 5 |
| Oligo F | SynLib_plex_n_f | 10 μM | 5 |
| Oligo R | SynLib_plex_n_r | 10 μM | 5 |
| Master Mix | NEBNext Ultra II Q5 Master Mix | 2x | 25 |
| Water |  |  | 10 |
|  |  | **Total volume** | 50 |
| **Time** | **Temperature (°C)** | **Number of cycles** |  |
| 30 sec | 98 | 1x |  |
| 15 sec | 98 | 18x |  |
| 30 sec | 68 |  |  |
| 40 sec | 72 |  |  |
| 2 min | 72 | 1x |  |
| Hold | 4 |  |  |

